# Transsulfuration links aspartate–asparagine metabolism and redox homeo-stasis to drive tumor growth

**DOI:** 10.64898/2026.07.10.737826

**Authors:** Mirko Milosevic, Kristina Dmytruk, Ahmad Alghadi, Pavel Jakoube, Julian Wong Soon, Petra Hyrossova, Muhammad Bin Munim, Sara Isabel Fernandes, Anna Shevzov-Zebrun, Robert Stanko, Isidora Mitric, Zuzana Cockova, Lukas Kucera, Juan Fernández-García, Ales Benda, Bryan Marzullo, Radislav Sedlacek, Jiri Neuzil, Sarah-Maria Fendt, Daniel A. Tennant, Matthew G. Vander Heiden, Katerina Rohlenova, Jakub Rohlena

**Author notes:** co-corresponding author <u>Editorial correspondence</u> or.

## Abstract

Cytosolic redox balance is tightly coupled to aspartate synthesis through the malate-aspartate shuttle, and limiting the malate-aspartate shuttle has been proposed to constrain tumor growth by restricting aspartate availability. Here we show that tumors derived from cancer cells lacking GOT1 and GOT2, the cytosolic and mitochondrial aspartate aminotransferases essential for aspartate production and malate-aspartate shuttle function, grow despite impaired canonical aspartate synthesis. This is because cytosolic redox state, not aspartate supply, is the primary metabolic bottleneck in GOT1/GOT2 knockout cells. Using single-cell transcriptomics, metabolite tracing, and a loss-of-function CRISPR screen, we find that these tumors engage an adaptive bypass in which availability of asparagine, a product of aspartate, enables serine- and methio-nine-dependent transsulfuration to generate α-ketobutyrate, whose reduction regenerates cytosolic NAD⁺ and restores redox homeostasis. Pharmacological inhibition or genetic ablation of transsulfuration abrogates this asparagine-driven rescue. These findings define asparagine as a regulator of cytosolic NAD⁺/NADH balance and reveal a link between amino acid metabolism and redox control that suggests transsulfuration as a targetable vulnerability in tumor redox maintenance.

**Significance statement:** Aspartate synthesis and cytosolic redox balance are both coupled through the malate-aspartate shuttle. We show that the cytosolic NAD⁺/NADH ratio, not aspartate supply, is a critical output of the malate-aspartate shuttle for tumor growth. Availability of asparagine, a product of aspartate, enables serine- and methionine-dependent transsulfuration to restore cytosolic NAD⁺/NADH balance, proliferation and tumor growth independently of canonical aspartate production by the malate-aspartate shuttle. This defines asparagine as a regulator of cytosolic redox and identifies transsulfuration as a targetable vulnerability in tumor redox maintenance.

## Introduction

Cancer cell proliferation depends on biosynthetic pathways that supply metabolic precursors for nucleotides, amino acids, and lipids. Under nutrient-limited or hypoxic conditions, specific pathways, often those that involve redox metabolism, can become selectively essential, presenting opportunities for therapeutic targeting. Among these, aspartate plays a central role as a precursor for asparagine, arginine, and purine/pyrimidine nucleotides^1–3^.

Aspartate can be synthesized by either of two aspartate (a.k.a glutamic oxaloacetic) aminotransferases: GOT1, located in the cytosol, and GOT2, located in mitochondria^4,5^. Both enzymes are integral components of the malate-aspartate shuttle (MAS), which helps maintain compartmentalized NAD⁺/NADH ratio by transferring reducing equivalents across the mitochondrial membrane for NAD⁺ regeneration by the respiratory chain. The MAS thus supports biosynthesis in two ways: (i) directly by producing aspartate and (ii) indirectly by maintaining cytosolic redox homeostasis. When mitochondrial respiration is impaired—such as under hypox-ia—reduced NAD⁺ availability stalls the MAS, disrupts cytosolic redox state, limits aspartate synthesis and constrains tumor growth^1,6–8^.

Previous studies have suggested that aspartate shortage can be a key bottleneck for proliferation in such redox-compromised contexts^1,7,8^. These studies also highlighted a central role for cytosolic redox in providing aspartate: restoring cytosolic NAD^+^ availability enables aspartate synthesis and rescues proliferation^6^. However, redox imbalance that precedes aspartate shortage can also independently impair biosynthesis^9,10^. Indeed, proliferating cells employ multiple mechanisms to maintain the cytosolic NAD+/NADH ratio, including the glycerol-3-phosphate shuttle and fatty acid desaturation^11,12^. Dissecting whether aspartate availability or cytosolic redox state is the primary growth-limiting factor is critical for understanding cancer metabolic flexibility and for guiding strategies to target metabolism for cancer therapy. Here, we dissect the importance of the MAS in cytosolic redox control versus aspartate production by generating cancer cell models lacking both GOT1 and GOT2, thereby simultaneously blocking canonical aspartate synthesis and MAS-mediated redox maintenance. Combined with single-cell transcriptomics, metabolic tracing, and a targeted CRISPR screen in cancer cells and tumors, this approach allowed us to disentangle the contributions of aspartate supply and cytosolic redox balance to supporting tumor growth in these models and identify an unexpected aspara-gine/transsulfuration bypass of the combined MAS /aspartate biosynthesis defect (Fig. 1A,B).

**Figure 1:**
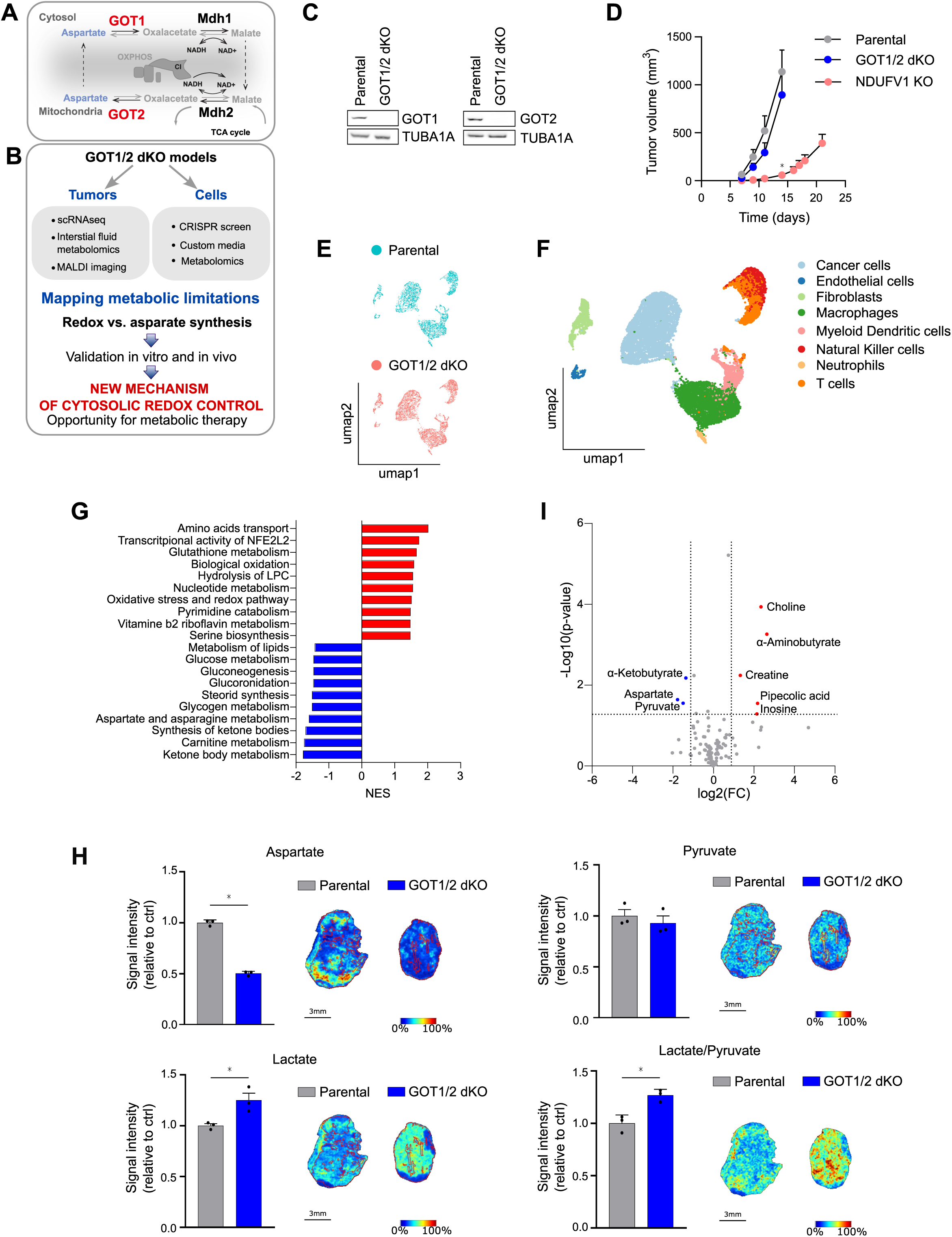
GOT1 and GOT2 deficient cells produce tumors normally. **A**) GOT1 and GOT2 transaminases in the context of the malate-aspartate shuttle. **B**) Design of the study. **C**) Deletion of GOT1 and GOT2 protein in GOT1/2 dKO (double knockout) B16 cells determined by western blot using antibodies against the N-termini. Representative data from multiple re-peated measurements. **D**) Tumor growth kinetics of mouse melanoma tumors generated by s.c. injection of parental, GOT1/2 dKO or NDUFV1 KO cells (mean ± SEM, n=6, *p < 0.05 compared to parental tumors, two-way ANOVA). Note: Statistics was calculated at day 14, when the parental and GOT1/2 dKO tumors were harvested due to reaching the ethical endpoint. **E**) UMAP dimensionality reduction plot of single cell transcriptome originating from parental or GOT1/2 dKO tumors, color-coded by sample. **F**) UMAP dimensionality reduction plot of single-cell transcriptome from parental and GOT1/2 dKO tumors, color-coded by cell type. **G**) Metabolic gene set enrichment analysis in GOT1/2 dKO versus parental cancer cell subset (p < 0.05). **H**) Intratumoral levels of aspartate, lactate and pyruvate, and the lactate/pyruvate ratio, in pa-rental and GOT1/2 dKO B16 melanoma tumors determined by MALDI imaging (mean ± SEM, n = 3, ∗p < 0.05, unpaired two-tailed t test). A representative image is shown on the right. **I**) Quantification of metabolites in the tumor interstitial fluid of GOT1/2 dKO versus parental tumors, measured by LC-MS. Blue indicates significantly decreased metabolites and red indi-cates significantly increased metabolites in GOT1/2 dKO tumors relative to parental B16 con-trols (n=3, significant hits log2FC > 1, p < 0.05).

## Results

### GOT1/2 DEFICIENT CANCER CELLS GENERATE TUMORS NORMALLY

GOT1 and GOT2 are enzymes that are important for both aspartate synthesis and cellular redox maintenance via the MAS. Previous reports showed that respiratory chain-deficient cells cannot form tumors^9,10,13,14^ or synthesize aspartate^6,7^. We hypothesized that similarly to defects in respiration, simultaneous ablation of GOT1 and GOT2 might severely impair tumor formation. To test this, we generated cancer cells with double knockout (dKO) of GOT1 and GOT2 by CRISPR-mediated gene editing in B16 melanoma, 4T1 mammary carcinoma, and HKP1 lung adenocarcinoma cells. Editing of the targeted loci was confirmed by PCR and the absence of the GOT1 and GOT2 protein by western blot using antibodies against the N- and C termini of the two transaminases (Fig. 1C, S1A-D). Unexpectedly, when injected into syngeneic mice, GOT1/2 ablated cancer cells produced tumors that grew at a similar rate as GOT1/2 intact controls (Fig. 1D and Fig. S1E, F). This result, consistent across models, suggests a robust compensatory mechanism that overcomes the defect of aspartate synthesis as well as the dysregulated cytosolic redox state from the loss of MAS. To directly assess how combined GOT1/2 deficiency compares with a respiratory chain defect in terms of tumor growth, we eliminated NDUFV1, the catalytic subunit of mitochondrial complex I. In contrast to GOT1/2 dKO, knockout of NDUFV1 strongly impaired tumor growth in the B16 and 4T1 models (Fig. 1D, S1E), consistent with prior reports^9,10,13,14^. This demonstrates that unlike a mitochondrial complex I defect, loss of endogenous aspartate synthesis and cytosolic redox maintenance is readily compensated to enable tumor growth. Therefore, suppressed aspartate synthesis may not necessarily contribute to the poor tumor-forming capacity of respiration-deficient cells^9,15,16^.

As GOT1/2 dKO cancer cells form tumors normally, we next aimed to elucidate how these cells adapt to GOT1/2 deficiency. We collected B16 parental and GOT1/2 dKO tumors and performed single-cell RNA sequencing (scRNA-seq, Fig. 1E). We identified all the major cell types expected to be found in a tumor microenvironment, including cancer cells, endothelial cells, fibroblasts and several subsets of immune cells (Fig.1F). While the absence of GOT1/2 in cancer cells did not substantially alter the fractional composition of tumor resident cell types (Fig. S1G), it impacted the transcriptome of both cancer and stromal cells. Focusing on metabolic genes, in the absence of GOT1 and GOT2 we found upregulation of amino acid transport / metabolism in macrophages, endothelial cells and cancer cells (Fig. S1H, Table S1). In cancer cells, amino acid transport was the most upregulated metabolic pathway (Fig. 1G, Table S1), suggesting that GOT1/2 dKO cells may rely on importing amino acids. Previously it was reported that environmental limitations such as hypoxia can stimulate aspartate salvage from the environment in K-RAS mutated pancreatic adenocarcinoma^17^. We thus further sub-clustered cancer cells to examine whether the upregulation of amino acid transport is specific to a particular cell state, such as to cells constrained by the lack of nutrients or oxygen. This refined analysis yielded 5 distinct clusters of cancer cells that could be identified by their uniquely assigned markers as inflammatory (cluster 0), proliferative (cluster 1), nutrient-challenged/hypoxic (cluster 2), proliferative/inflammatory (cluster 3) and transitioning (cluster 4) (Fig. S1I, J). Amino acid transport was increased in four out of five clusters, while the fifth cluster (cluster 4) consistently showed upregulation of amino acid metabolism (Fig. S1K, Table S2). This points to a strong impact of GOT1/2 deficiency on amino-acid homeostasis in cancer cells in vivo, independent of the cancer cell state. Overall, the growth rate of GOT1/2 deficient cancer cells in vivo might be, at least in part, supported by stroma-produced nutrients, in particular amino acids.

To directly probe the metabolic impact of GOT1/2 deletion in vivo, we employed MALDI imaging to visualize aspartate levels and the lactate/pyruvate ratio as an inverse proxy for cytosolic NAD^+^/NADH. Despite little impact on the growth rate, tumors arising from GOT1/2 dKO cells featured reduced aspartate levels and a lower pyruvate/lactate ratio (Fig. 1H), the expected phenotype of MAS deficiency. To test whether GOT1/2 deficient cells sustain proliferation in vivo by compensatory uptake of nutrients from the tumor microenvironment, we quantified absolute levels of 107 polar metabolites in tumor interstitial fluid. This analysis revealed a marked depletion of aspartate, pyruvate and α-ketobutyrate in the interstitial fluid of GOT1/2 dKO compared to parental tumors (Fig. 1I, Table S3), while the plasma concentration of these metabolites was not different (Fig. S1L, Table S4). These results reinforce the notion that GOT1/2 dKO cancer cells in tumors may rely on environmental nutrients to sustain their growth and likely undergo metabolic rewiring that engages specific metabolic pathways to compensate for disruption of GOT1/2.

#### Exogenous pyruvate and α-ketobutyrate but not aspartate support proliferation of GOT1/2 DEFICIENT CELLS

To understand the metabolic limitations that the GOT1/2-derived tumors must overcome to proliferate, we used an array of commercial and custom-made growth media where we supplemented or removed individual nutrients in vitro. We focused on amino acids (based on scRNAseq results) and on aspartate, pyruvate and α-ketobutyrate, metabolites depleted in the interstitial fluid of GOT1/2 dKO tumors (c.f. Fig. 1I). First, we assessed proliferation in RPMI and DMEM, two frequently used standard culture media that differ in the availability of amino acids. RPMI has more amino acids than DMEM and contains a similar concentration of aspartate to that found in the interstitial fluid of GOT1/2 dKO tumors (153.1 +/- 41.09 µM, SEM), whereas DMEM has fewer amino acids and has no aspartate. In RPMI medium the GOT1/2 dKO B16, 4T1 and HKP1 cells proliferated similarly to parental cells, recapitulating the in vivo findings. In contrast, all GOT1/2 dKO cells failed to proliferate in DMEM medium (Fig. 2A, Fig. S2 A,B).

**Figure 2:**
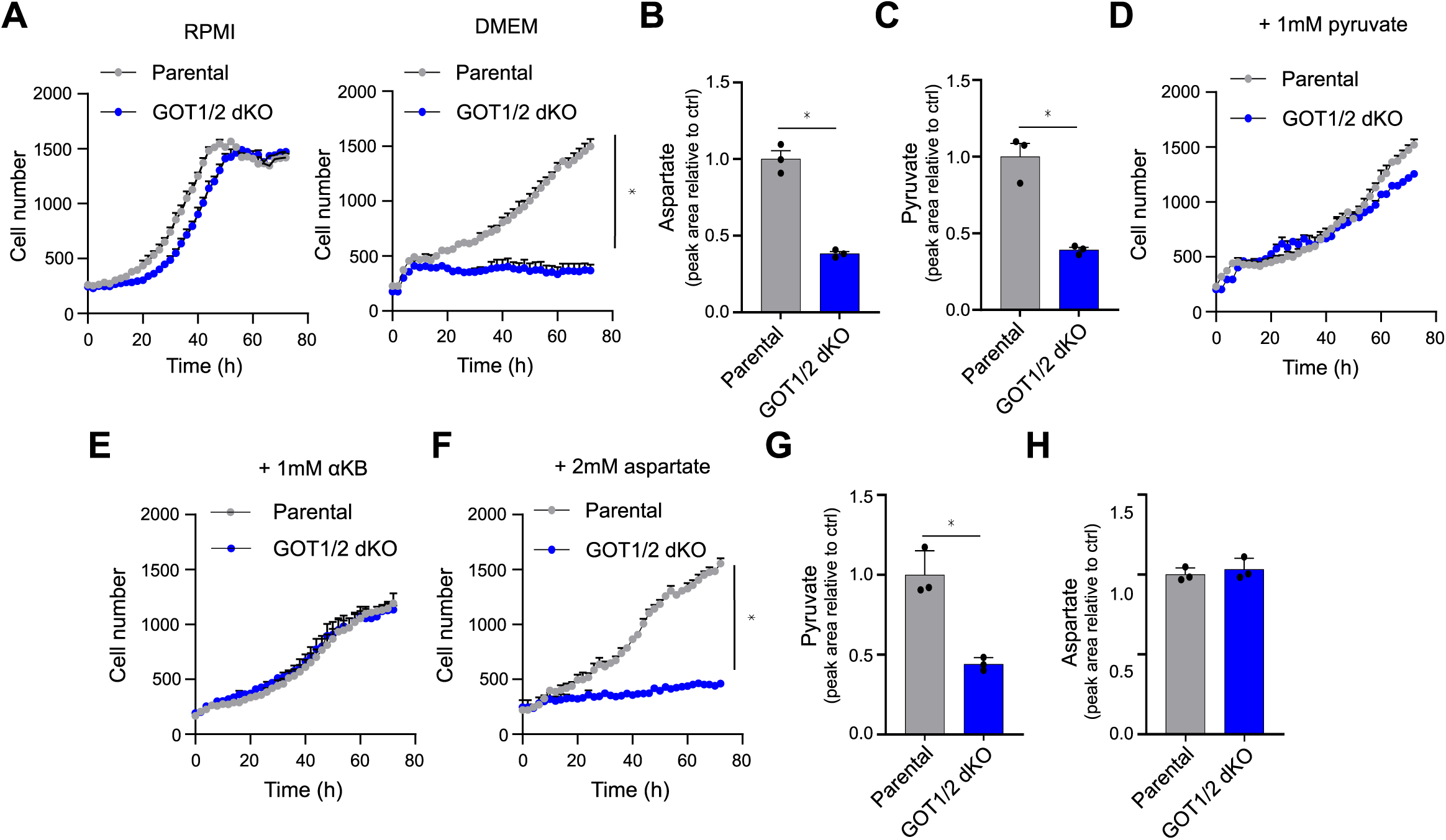
Context-dependent proliferation of GOT1 and GOT2 deficient cells. **A**) Proliferation of parental and GOT1/2 dKO cells in RPMI (left) and DMEM (right), measured by real-time cell proliferation assay (mean ± SEM, n = 3, ∗p < 0.05, unpaired two-tailed t test). **B**) Intracellular levels of aspartate in parental and GOT1/2 dKO cultured in DMEM medium, measured by GC-MS (mean ± SEM, n = 3, ∗p < 0.05, unpaired two-tailed t test). **C**) Intracellular levels of pyruvate in parental and GOT1/2 dKO grown in DMEM medium, meas-ured by GC-MS (mean ± SEM, n = 3, ∗p < 0.05, unpaired two-tailed t test). **D**) Proliferation of parental and GOT1/2 dKO cells in DMEM medium, supplemented with 1 mM pyruvate, measured by real-time cell proliferation assay (mean ± SEM, n = 3, unpaired two-tailed t test). **E**) Proliferation of parental and GOT1/2 dKO cells in DMEM medium, supplemented with 1 mM αKB, measured by real-time cell proliferation assay (mean ± SEM, n = 3, unpaired two-tailed t test). **F**) Proliferation of parental and GOT1/2 dKO cells in DMEM medium supplemented with 2 mM aspartate, measured by real-time cell proliferation assay (mean ± SEM, n = 3, ∗p < 0.05, un-paired two-tailed t test). **G**) Extracellular levels of pyruvate in medium collected from parental and GOT1/2 dKO cells, grown in RPMI supplemented with 1 mM pyruvate, measured by GC-MS (mean ± SEM, n = 3, ∗p < 0.05, unpaired two-tailed t test). **H**) Extracellular levels of aspartate in medium collected from parental and GOT1/2 dKO cells, grown in RPMI supplemented with 1 mM pyruvate measured by GC-MS (mean ± SEM, n = 3, ∗p < 0.05, unpaired two-tailed t test).

We next measured intracellular levels of aspartate and pyruvate (by GC-MS which did not measure α-ketobutyrate) in DMEM-cultured GOT1/2 dKO B16 cells. Both aspartate and pyruvate were depleted (Fig. 2B, C), suggesting that reduced levels of these metabolites in the interstitial fluid of GOT1/2 dKO tumors (c.f. Fig. 1I) could result from compensatory uptake that maintains proliferation. To test this premise directly, we supplemented DMEM medium with aspartate, pyruvate or α-ketobutyrate. Whereas pyruvate or α-ketobutyrate supplementation fully restored proliferation of GOT1/2 dKO B16 cells in DMEM, (Fig. 2D, E), aspartate had no effect (Fig. 2F). These findings were confirmed in 4T1 and HKP1 cell lines (Fig.S2 C, D). Hence, although both intracellular pyruvate and aspartate are depleted in GOT1/2 dKO cells and in tumor interstitial fluid, only pyruvate (and α-ketobutyrate) supplementation rescues proliferation. Consistently, pyruvate, but not aspartate, was depleted from pyruvate and aspartate-containing culture medium (Fig. 2G, H). In RPMI where GOT1/2 dKO proliferate normally, supplementation with pyruvate, α-ketobutyrate or aspartate did not further increase proliferation (Fig. S2E-F), indicating that nutrients present in RPMI eliminated the dependence of GOT1/2 cells on pyruvate or α-ketobutyrate supplementation. To summarize, extracellular pyruvate and α-ketobutyrate support proliferation of GOT1/2 dKO in DMEM, but extracellular aspartate does not. This suggests that in GOT1/2 tumors pyruvate and α-ketobutyrate are likely depleted from the extracellular environment because of increased consumption, whereas the reduced level of aspartate might reflect its decreased production by cancer cells.

### Aspartate availability does not limit GOT1/2 dKO cells

Because aspartate supplementation did not rescue proliferation of GOT1/2 dKO cells in DMEM medium (cf. Fig. 2F, S2 C, D), we investigated how these cells obtain aspartate. Specifically, we tested whether GOT1/2 dKO cells can take up exogenous aspartate and if they retain an ability to synthesize aspartate endogenously. The cell membrane is poorly permeable for aspartate, and uptake is thus largely transporter-mediated^8^. SLC1A3, the main transporter for aspartate, was not upregulated in GOT1/2 dKO cells (Fig S2G). Further, the SLC1A3 inhibitor TFB-TBOA did not reduce proliferation in aspartate-containing RPMI (Fig. S2H). This suggests that any SLC1A3 expressed by GOT1/2 dKO cells might not be functionally relevant and that free extracellular aspartate is not taken up by GOT1/2 dKO cells. To test this directly, we incubated cells with uniformly labeled U-^13^C aspartate (U-^13^C Aspartate) and measured the amount of U-^13^C aspartate in the cells and in the media. Consistently, U-^13^C aspartate was not detected in cells and was not depleted from the medium (Fig. S2I), confirming that GOT1/2 absence in these cells is not compensated for by uptake of free environmental aspartate.

Another route to obtain aspartate from the extracellular space, reported in RAS mutant cells, is macropinocytosis of intact extracellular proteins^17^. We tested utilization of macropinocytosis in B16 cancer cell model, which are RAS wild type^18^. Macropinocytotic index was not increased in GOT1/2 dKO cells (Fig S2J), and a macropinocytosis inhibitor EIPA did not reduce proliferation rate of GOT1/2 dKO cells in RPMI (Fig. S2K, c.f. Fig. 2A). Likewise, supplementation of 2% BSA, a substrate for macropinocytosis, did not rescue proliferation of GOT1/2 dKO cells in DMEM (Fig S2L), where proliferation is restricted (c.f. Fig. 2A). Hence, GOT1/2 dKO cells do not appear to use macropinocytosis to overcome GOT1/2 loss and likely obtain aspartate via a route other than catabolism of extracellular protein.

We next used metabolic tracing to assess endogenous aspartate synthesis in GOT1/2 dKO cells. Despite a lack of GOT1 and GOT2 (cf. Fig. 1C, S1A), tracing from U-^13^C labeled glucose or glutamine revealed a low level of continued de novo synthesis (Fig. S2M, N). This suggests that an alternative pathway(s) must enable a limited amount of aspartate synthesis in a GOT1/2-independent manner, which is sufficient to meet aspartate needs in the absence of GOT1/2. This is in line with a recent report showing that proliferation is still supported even when intracellular aspartate levels are substantially reduced^19^. This is also consistent with factors other than aspartate availability limiting the proliferation of GOT1/2 dKO cells.

### A LOW NAD^+^ /NADH RATIO IN THE CYTOSOL IS A LIMITATION FOR GOT1/2 DKO CELL PROLIFERATION

GOT1/2 dKO-induced disruption of MAS is expected to alter cytosolic redox balance. Pyruvate and α-ketobutyrate are exogenous electron acceptors that can regenerate NAD^+^ from NADH in a reaction catalyzed by cytosolic dehydrogenases such as lactate dehydrogenase^7,17,20^. Their depletion in the interstitial fluid of GOT1/2 dKO tumors (c.f. Fig. 1I) might reflect their increased uptake to restore redox balance. Therefore, redox, not aspartate availability, may limit proliferation of GOT1/2 dKO cells. We first confirmed that GOT1/2 dKO cells exhibit a low NAD^+^/NADH ratio (Fig. 3A) due to cytosolic NADH accumulation (Fig. 3B). We next used pyruvate to test if recovering the redox state promotes GOT1/2-independent proliferation. Consistent with py-ruvate-mediated rescue of GOT1/2 dKO cells in DMEM (c.f. Fig. 2D), the cytosolic NAD^+^/NADH ratio was normalized by pyruvate supplementation in these conditions (Fig. 3C, S3A), and pharmacological inhibition of pyruvate uptake (and lactate secretion) suppressed proliferation of GOT1/2 dKO cells and the growth of GOT1/2 dKO tumors (Fig. 3D, E). Exogenous pyruvate can thus at least partially bypass the GOT1/2 defect and restore proliferation both in vitro and in vivo.

**Figure 3:**
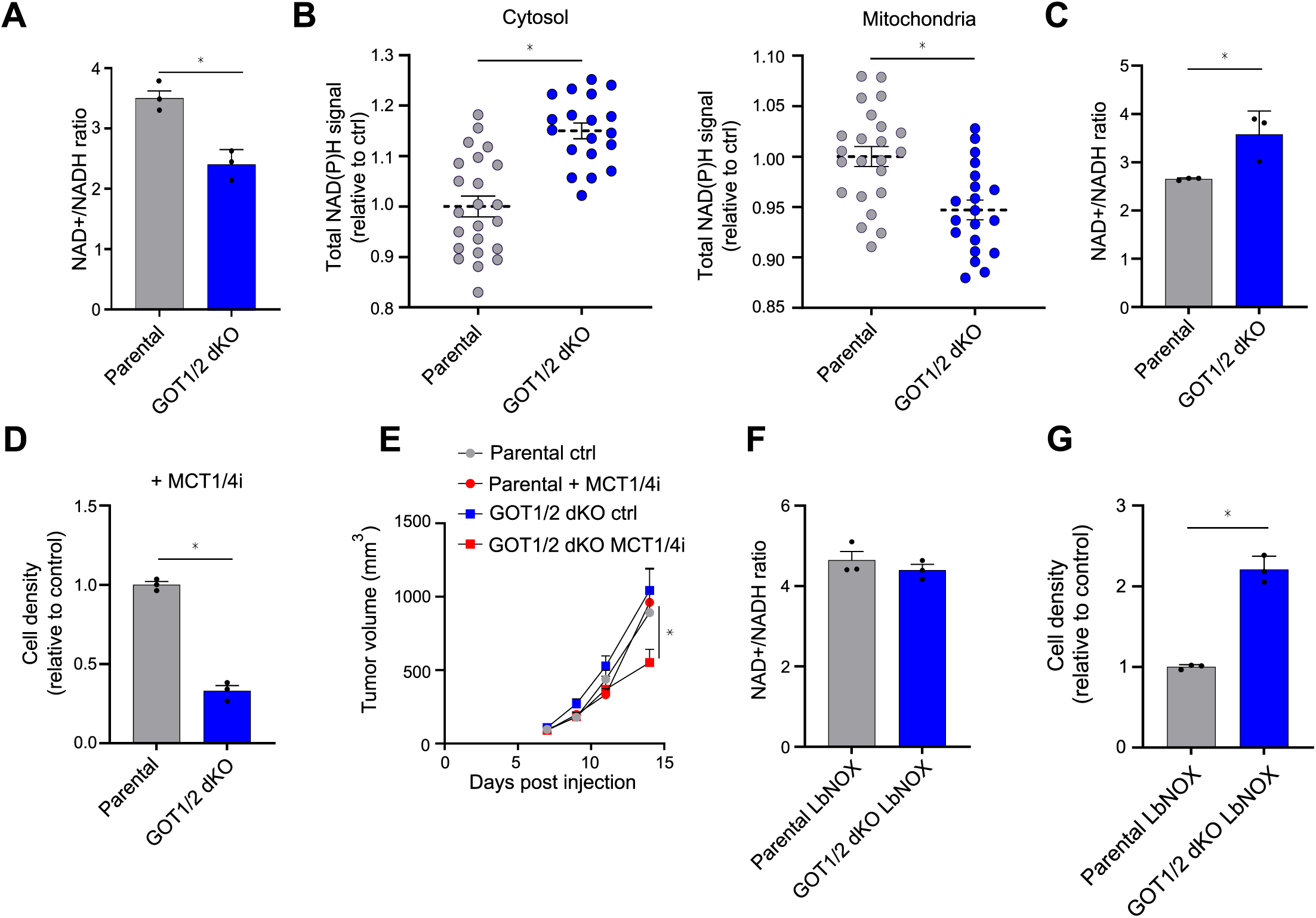
Combined GOT1 and GOT2 deficiency disrupts cytosolic redox. **A**) NAD+/NADH ratio in parental and GOT1/2 dKO cells grown in DMEM medium, measured by bioluminescent assay (mean ± SEM, n = 3, ∗p < 0.05, unpaired two-tailed t test). **B**) Total NAD(P)H signal in the cytosol and in mitochondria of parental and GOT1/2 dKO cultured in DMEM medium, measured by 2-photon fluorescence-lifetime imaging microscopy (mean ± SEM, n > 19 wells, ∗p < 0.05, unpaired two-tailed t test). **C**) NAD+/NADH ratio in parental and GOT1/2 dKO cells grown in DMEM medium, in the pres-ence of 1 mM pyruvate, measured by luminescent enzymatic assay (mean ± SEM, n = 3, ∗p < 0.05, unpaired two-tailed t test). **D**) Proliferation of parental and GOT1/2 dKO cells in the presence of 20µM MCT1/4 inhibitor grown DMEM medium, supplemented with 1mM pyruvate, measured by crystal violet assay after 72 h (mean ± SEM, n = 3, ∗p < 0.05, unpaired two-tailed t test). **E**) Tumor growth kinetics of mouse melanoma tumors generated by s.c. injection of parental and GOT1/2 dKO cells and treated with syrosingopine (MCT1/4 inhibitor) (5.5 mg/kg of mouse) or with vehicle (corn oil) every 24 h (mean ± SEM, n = 5, ∗p < 0.05, two-way ANOVA). **F**) NAD+/NADH ratio in parental and GOT1/2 dKO cells grown in DMEM medium, with expres-sion of cytosolic LbNOX, measured by bioluminescence assay (mean ± SEM, n = 3, ∗p < 0.05, unpaired two-tailed t test). **G**) Proliferation of parental and GOT1/2 dKO cells in DMEM medium, with expression of cyto-solic LbNOX, measured by crystal violet assay after 72 h (mean ± SEM, n = 3, ∗p < 0.05, unpaired two-tailed t test). Note that Lb-NOX expression reduced proliferation of parental cells.

Because mitochondrial oxidation potentially represents a major sink for cytosolic pyruvate, we tested whether altering mitochondrial pyruvate utilization affects proliferation of GOT1/2 dKO cells. A pyruvate dehydrogenase kinase inhibitor, which increases utilization of pyruvate in mitochondria, limited proliferation of GOT1/2 dKO cells (Fig. S3B). Conversely, an inhibitor of mitochondrial pyruvate carrier which restricts pyruvate import into mitochondria, conferred a relative advantage to GOT1/2 dKO cells compared to parental cells (Fig. S3C). This suggests that cytosolic, but not mitochondrial pyruvate, is required to compensate for the GOT1/2 loss, likely by restoring cytosolic redox state.

To directly assess whether cytosolic redox restoration rescues proliferation of GOT1/2 dKO cells, we used LbNOX, an alternative oxidase that can selectively elevate the NAD^+^/NADH ratio in a compartment-specific manner independent of pyruvate (Fig. S3D)^21^. Consistent with impaired transfer of reducing equivalents from the cytosol to mitochondria in the absence of GOT1 and GOT2, expression of LbNOX in the mitochondria had little or no effect on either the cellular NAD^+^/NADH ratio or proliferation (Fig. S3E-F). In contrast, expression of LbNOX in the cytosol normalized the NAD^+^/NADH ratio (Fig. 3F) and fully restored proliferation of GOT1/2 dKO cells (Fig. 3G). These findings confirm that disruption of cytosolic redox homeostasis is the main metabolic bottleneck that limits proliferation of GOT1/2 dKO cells, and that redox normalization underlies the rescue of proliferation by pyruvate.

### Asparagine, methionine and serine restore cytosolic redox to enable proliferation of GOT1/2 DKO CELLS

Having established that GOT1/2 dKO proliferation is limited by a low cytosolic NAD^+^/NADH ratio, we asked how these cells proliferate normally in RPMI (cf. Fig. 2A) even in the absence of pyruvate supplementation or LbNOX expression. We reasoned that RPMI-specific nutrients must normalize the cytosolic redox state another way to enable proliferation in the absence of GOT1 and GOT2. Indeed, the low NAD^+^/NADH ratio observed in DMEM-cultured GOT1/2 dKO cells was partially restored in RPMI (Fig. 4A). To identify nutrients that enable proliferation of GOT1/2 dKO cells, we customized RPMI medium by removing amino acids, individually or in subsets. Removal of most amino acids, including aspartate, had no or minimal impact on the proliferation rate of GOT1/2 dKO cells compared to parental cells in RPMI (Fig. S4A). In contrast, removal of methionine, serine or asparagine selectively suppressed GOT1/2 dKO proliferation in RPMI but did not affect the proliferation rate of parental cells (Fig. 4B-D). Pyruvate supplementation or LbNOX expression rescued proliferation of GOT1/2 dKO cell even in the absence of any of these three amino acids (Fig. S4B, C). To confirm these findings, we asked if supplementing DMEM, where GOT1/2 cells do not proliferate without redox normalization, with these amino acids could rescue proliferation. Because DMEM already contains methionine and serine, we added asparagine to RPMI levels. Asparagine supplementation elevated the NAD^+^/NADH ratio and fully restored proliferation of GOT1/2 dKO B16 cells (Fig. 4E, F), which was confirmed for 4T1 and HKP1 cells with GOT1/2 dKO (Fig. S4 D, E). Together, these results point to a causal and non-redundant role for methionine, serine and asparagine in cytosolic redox maintenance.

**Figure 4:**
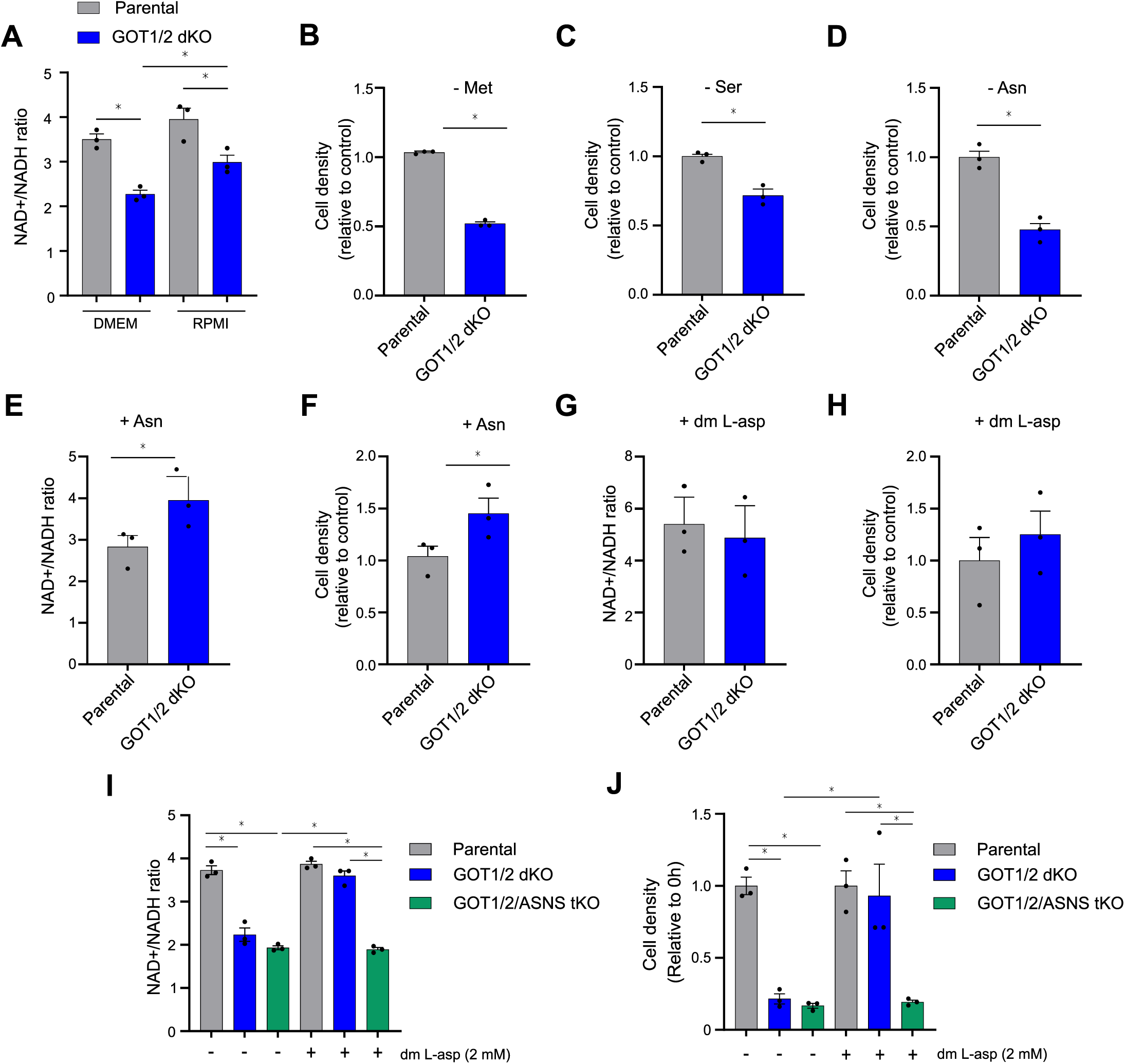
Asparagine metabolism restores redox balance in GOT1/2 dKO cells. **A**) NAD+/NADH ratio in parental and GOT1/2 dKO cells in DMEM and RPMI medium, measured by bioluminescence assay (mean ± SEM, n = 3, ∗p < 0.05, unpaired two-tailed t test). The DMEM data are the same as in panel 3A. **B**) Proliferation of parental and GOT1/2 dKO cells in RPMI medium, in the presence/absence of methionine (100 µM) measured by crystal violet assay after 72 h (mean ± SEM, n = 3, ∗p < 0.05, unpaired two-tailed t test). **C**) Proliferation of parental and GOT1/2 dKO cells in RPMI medium, in the presence/absence of serine (285 µM) measured by crystal violet assay after 72 h (mean ± SEM, n = 3, ∗p < 0.05, un-paired two-tailed t test). **D**) Proliferation of parental and GOT1/2 dKO cells in RPMI medium, in the presence/absence of asparagine (340 µM) measured by crystal violet assay after 72 h (mean ± SEM, n = 3, ∗p < 0.05, unpaired two-tailed t test). **E**) NAD+/NADH ratio in parental and GOT1/2 dKO cells in DMEM medium, in the ab-sence/presence of asparagine (340 µM) measured by bioluminescence omega assay (mean ± SEM, n = 3, ∗p < 0.05, unpaired two-tailed t test). **F**) Proliferation of parental and GOT1/2 dKO cells in DMEM medium, in the absence/presence of asparagine (340 µM) measured by crystal violet assay after 72 h (mean ± SEM, n = 3, ∗p < 0.05, unpaired two-tailed t test). **G**) NAD+/NADH ratio in parental and GOT1/2 dKO cells in DMEM medium, in the ab-sence/presence of dimethyl L-aspartate (2 mM) measured by bioluminescence assay (mean ± SEM, n = 3, ∗p < 0.05, unpaired two-tailed t test). **H**) Proliferation of parental and GOT1/2 dKO cells in DMEM medium, in the absence/presence of dimethyl L-aspartate (2 mM) measured by crystal violet assay after 72 h (mean ± SEM, n = 3, ∗p < 0.05, unpaired two-tailed t test). **I**) NAD+/NADH ratio in parental, GOT1/2 dKO and GOT1/2/ASNS tKO cells in DMEM medium, in the absence/presence of dimethyl L-aspartate (2 mM) measured by bioluminescence assay (mean ± SEM, n = 3, ∗p < 0.05, unpaired two-tailed t test). **J**) Proliferation of parental, GOT1/2 dKO and GOT1/2/ASNS tKO cells in DMEM medium, in the absence/presence of dimethyl L-aspartate (2 mM) measured by crystal violet assay after 72 h (mean ± SEM, n = 3, ∗p < 0.05, unpaired two-tailed t test).

Redox normalization by asparagine has not previously been reported. Asparagine, unlike methionine and serine, is directly produced from aspartate in a reaction catalyzed by the enzyme asparagine synthase (ASNS). Endogenous synthesis of asparagine via ASNS is reported to be a major consumer of intracellular aspartate^16,19^, and aspartate can recover cytosolic NAD+/NADH ratio via the GOT1-MDH1 reactions of the MAS.^22^ Even though our model lacks GOT1 which blocks this route, the unexpected effect of asparagine supplementation on redox normalization could be secondary to depletion of the aspartate pool. Reduced intracellular asparagine levels in DMEM-cultured GOT1/2 dKO cells were normalized in asparagine-containing RPMI (Fig. S4F), consistent with ASNS being bypassed in a way that alleviates aspartate consumption. To interrogate whether asparagine supplementation corrects the NAD^+^/NADH ratio independently of the aspartate pool, we used dimethyl aspartate, a cell permeable derivative of aspartate. Dimethyl aspartate supplementation elevated the NAD^+^/NADH ratio and restored proliferation of GOT1/2 dKO cells in DMEM medium (Fig. 4G, H) which does not contain aspara-gine. In contrast, when ASNS was ablated by CRISPR/Cas9 in GOT1/2 dKO cells, dimethyl aspartate supplementation could rescue neither the NAD^+^/NADH ratio nor proliferation (Fig. 4I, J). Hence, to restore the cellular redox state and enable proliferation in the absence of GOT1 and GOT2, aspartate needs to yield asparagine, making asparagine a causative agent in this scenario, and excluding an indirect effect of reduced intracellular availability of aspartate. Collectively, these results suggest that asparagine, either supplied externally or synthesized de novo from aspartate, is important for normalizing the cellular redox state and can restore proliferation of GOT1/2 dKO cells in the presence of serine and methionine independent of pyruvate.

### A CRISPR SCREEN REVEALS DEPENDENCE ON TRANSSULFURATION IN THE ABSENCE OF GOT1/2

To identify genes that enable asparagine-induced proliferation of GOT1/2 dKO cells in RPMI in the absence of pyruvate, we performed a targeted loss of function CRISPR screen using an sgRNA library of 2865 mouse metabolic genes^23^. Parental and GOT1/2 dKO cells were grown in RPMI medium for 14 population doublings in the presence and absence of pyruvate after which the sgRNA repertoire was profiled by next generation sequencing (Fig. 5A). sgRNAs recovered at the end of the screen were compared to the starting population for each condition. Analysis suggested 20 (unique) genes selectively essential in the absence of GOT1 and GOT2 without pyruvate addition (Fig. 5B), but only 7 (unique) essential genes with pyruvate supple-mented in the medium (Table S5). The reduced number of essential genes in the presence of pyruvate is consistent with redox being a major limitation in GOT1/2 dKO cells. For further analysis, we first assigned the identified GOT1/2-specific essential genes to metabolic pathways and prioritized pathways that contained multiple hits. Multiple essential genes were identified in pathways that contribute to redox maintenance, including the mitochondrial respiratory chain. Interestingly, several hits pointed to the importance of transsulfuration and methionine/serine metabolism. These hits included SLC3A2, which is an essential subunit of the neutral amino acid transporter that imports methionine into the cell, and the cystine-glutamate antiporter, and cystathionine β-synthase (CBS), the first and rate limiting enzyme of the transsulfuration pathway ^24^ (Fig 5C). These findings, consistent with the auxotrophy of GOT1/2 dKO cells for methionine and serine (c.f. Fig. 4B, C), suggest that the transsulfuration pathway may enable aspara-gine-dependent proliferation. Hence, we focused next on uncovering how asparagine supports transsulfuration, and how transsulfuration promotes proliferation of GOT1/2 dKO cells.

**Figure 5:**
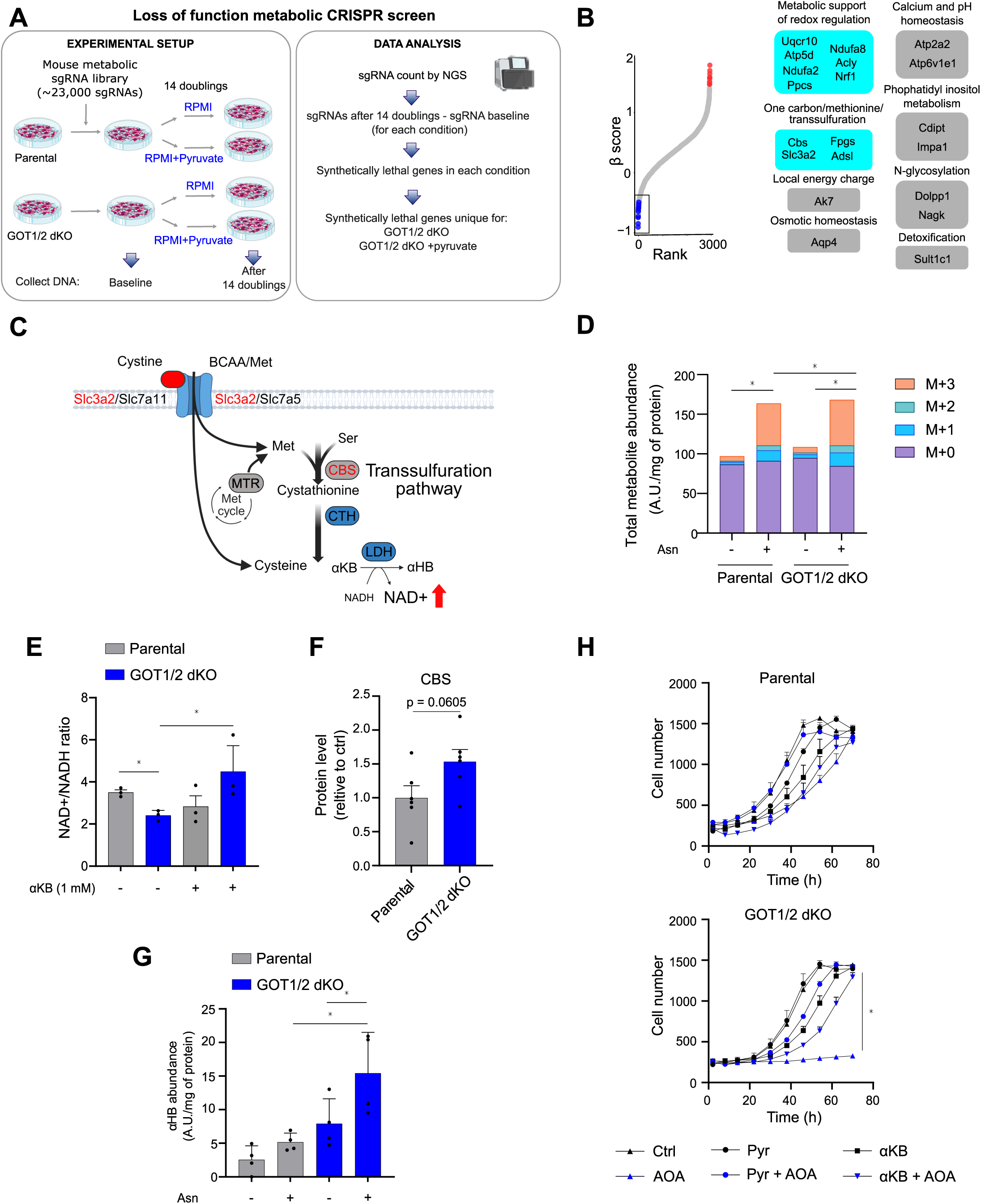
Crispr screen for genes that support GOT1/2 deficient cells. **A**) Scheme of experimental approach **B**) Waterfall plot of β scores from unique CRISPR screen hits, ranked and further subdivided based on their biological roles (blue significant negative selection hits, red significant positive selection hits, FDR < 0.25) **C**) Scheme of the Transsulfuration pathway and the related metabolism. CRISPR screen hits are highlighted in red. **D**) Total intracellular serine levels and isotopologue distribution in B16 cells preloaded or not with asparagine (340 µM) for 2h, followed by 2h incubation with U- C3-Serine in RPMI medium in the absence of cystine, methionine and asparagine, measured by LC-MS (mean ± SEM, n = 4, ∗p < 0.05, unpaired two-tailed t test calculated on the M+3 serine isotopologue). **E**) NAD+/NADH ratio in parental and GOT1/2 dKO cells in DMEM medium, in the ab-sence/presence of α-ketobutyrate (α-KB, 1 mM) measured by bioluminescence assay (mean ± SEM, n = 3, ∗p < 0.05, unpaired two-tailed t test). **F**) Quantification of CBS protein in parental and GOT1/2 dKO B16 cells determined by western blot (mean ± SEM, n = 5, unpaired two-tailed t test). **G**) Intracellular levels of α-hydroxybutyrate in parental and GOT1/2 dKO B16 cells grown in cus-tom RPMI medium in the absence of cystine and presence/absence of asparagine (340 µM) for 24 hours, measured by LC-MS (mean ± SEM, n = 4, ∗p < 0.05, unpaired two-tailed t test). **H**) Proliferation of parental and GOT1/2 dKO cells in RPMI medium, in the presence/absence of AOA (300 µM) supplemented with or without pyruvate (1 mM), α-ketobutyrate (α-KB, 1 mM), measured by real-time cell proliferation assay (mean ± SEM, n = 3, ∗p < 0.05, unpaired two-tailed t test).

### Asparagine promotes transsulfuration to support redox normalization and proliferation

Methionine and serine are transsulfuration substrates, but the link between transsulfuration and asparagine is less obvious. Asparagine has been reported to act as an exchange factor for uptake of extracellular serine^25^, raising the possibility that asparagine could support GOT1/2 dKO cells indirectly by facilitating serine import. To test this, we cultured parental and GOT1/2 dKO B16 cells in medium supplemented with U-^13^C-serine after preloading or not with asparagine and measured fractional U-^13^C enrichment in intracellular serine using LC-MS. Asparagine preloading increased both intracellular serine concentration and the abundance of the M+3 isotopologue, consistent with an asparagine-dependent increase in serine uptake.

Furthermore, the M+3 isotopologue of serine was enriched in asparagine-preloaded GOT1/2 dKO compared to asparagine-preloaded parental cells (Fig. 5D). This is consistent with increased expression of SLC1A5/ASCT2 (Fig. S4G), the asparagine/serine exchanger, in GOT1/2 dKO cells. These findings suggest that asparagine supports transsulfuration in GOT1/2 dKO cells at least in part by facilitating serine import.

CBS is the first enzyme of the transsulfuration pathway (Fig. 5C). It condenses serine with homocysteine, derived from methionine, to produce cystathionine. Cystathionine is then converted into cysteine, the final product, by the second enzyme of the pathway, cystathionine γ-lyase (CSE). In addition, the transsulfuration pathway is also a major producer of hydrogen sulfide. While cysteine and hydrogen sulfide are considered the main products, α-ketobutyrate is generated as a byproduct^24,26,27^. α-ketobutyrate, similarly to pyruvate, can normalize the cytosolic redox state via dehydrogenases that convert α-ketobutyrate into α-hydroxybutyrate while regenerating cytosolic NAD^+^ (Fig. 5C)^7,28^. In our model, supplementation with α-ketobutyrate, but not cystine or hydrogen sulfide, rescued proliferation of GOT1/2 dKO cells in DMEM medium (cf. Fig. 2 A,E, Fig. S5 A) and normalized their NAD^+^/NADH ratio (Fig. 5E), identifying α-ketobutyrate as the transsulfuration product responsible for GOT1/2 dKO cell rescue. Consistently, the CBS protein trended towards increased (Fig. 5F), and intracellular levels of α-hydroxybutyrate were markedly elevated in GOT1/2 dKO cells in the presence of asparagine (Fig. 5G). GOT1/2 dKO cells thus both take up more exogenous α-ketobutyrate (cf. Fig. 1I) and increase its endogenous production in an asparagine-dependent manner to normalize the cytosolic redox state by α-ketobutyrate to α-hydroxybutyrate conversion.

Next, we assessed whether disruption of transsulfuration selectively impairs GOT1/2 dKO cells. Both CBS and CSE rely on pyridoxal phosphate as a cofactor, which renders the transsulfuration pathway sensitive to a pyridoxal phosphate inhibitor aminooxyacetic acid (AOA). While AOA inhibits many pyridoxal phosphate-containing enzymes^29^, the transsulfuration pathway and transaminases participating in aspartate production are among its most prominent targets^30^. Strikingly, proliferation of GOT1/2 dKO B16 cells was selectively suppressed by AOA (Fig. 5H), which was confirmed in 4T1 and HKP1 models (Fig. S5B, C). The effect of AOA in these experiments was not mediated by inhibition of any (alternative) aspartate synthesis pathway, as the AOA concentration used did not reduce aspartate synthesis in parental or in GOT1/2 dKO cells (Fig. S5D). In RPMI, which contains asparagine, serine and methionine, proliferation and the NAD^+^/NADH ratio of AOA-treated GOT1/2 dKO B16, 4T1 and HKP1 cells were restored by supplementation with pyruvate or α-ketobutyrate, a product of transsulfuration, consistent with AOA targeting cellular redox regulation, not aspartate synthesis (Fig. 5H, S5E). In DMEM medium, which contains serine and methionine but lacks asparagine, supplementation with asparagine or dimethyl aspartate did not rescue the NAD^+^/NADH ratio or proliferation of AOA-treated GOT1/2 dKO cells (Fig. S5F-G). Collectively, these results suggest that aspartate and asparagine act upstream of transsulfuration.

To test if the transsulfuration pathway is essential for redox maintenance and proliferation using a genetic model, we knocked out CBS in both parental and GOT1/2 dKO cells, producing a CBS knockout (CBS KO) and a GOT1/2/CBS triple knockout (GOT1/2/CBS tKO) (Fig. 6A). Both CBS KO and GOT1/2/CBS tKO were sensitive to cystine removal (Fig. S6A, B), confirming that transsulfuration is disabled. In RPMI medium, which contains asparagine, the NAD^+^/NADH ratio of GOT1/2/CBS tKO cells was reduced compared to GOT1/2 dKO cells (Fig. 6B), in line with transsulfuration contributing to cytosolic redox maintenance. Proliferation of GOT1/2/CBS tKO cells in RPMI was impaired, whereas GOT1/2 dKO and CBS KO cells proliferated normally (Fig. 6C). Both redox and proliferation of GOT1/2/CBS tKO cells in RPMI were restored by pyruvate or α-ketobutyrate (Fig. 6D-G), but not by hydrogen sulfide or cystine supplementation (Fig. S6A-C).

**Figure 6:**
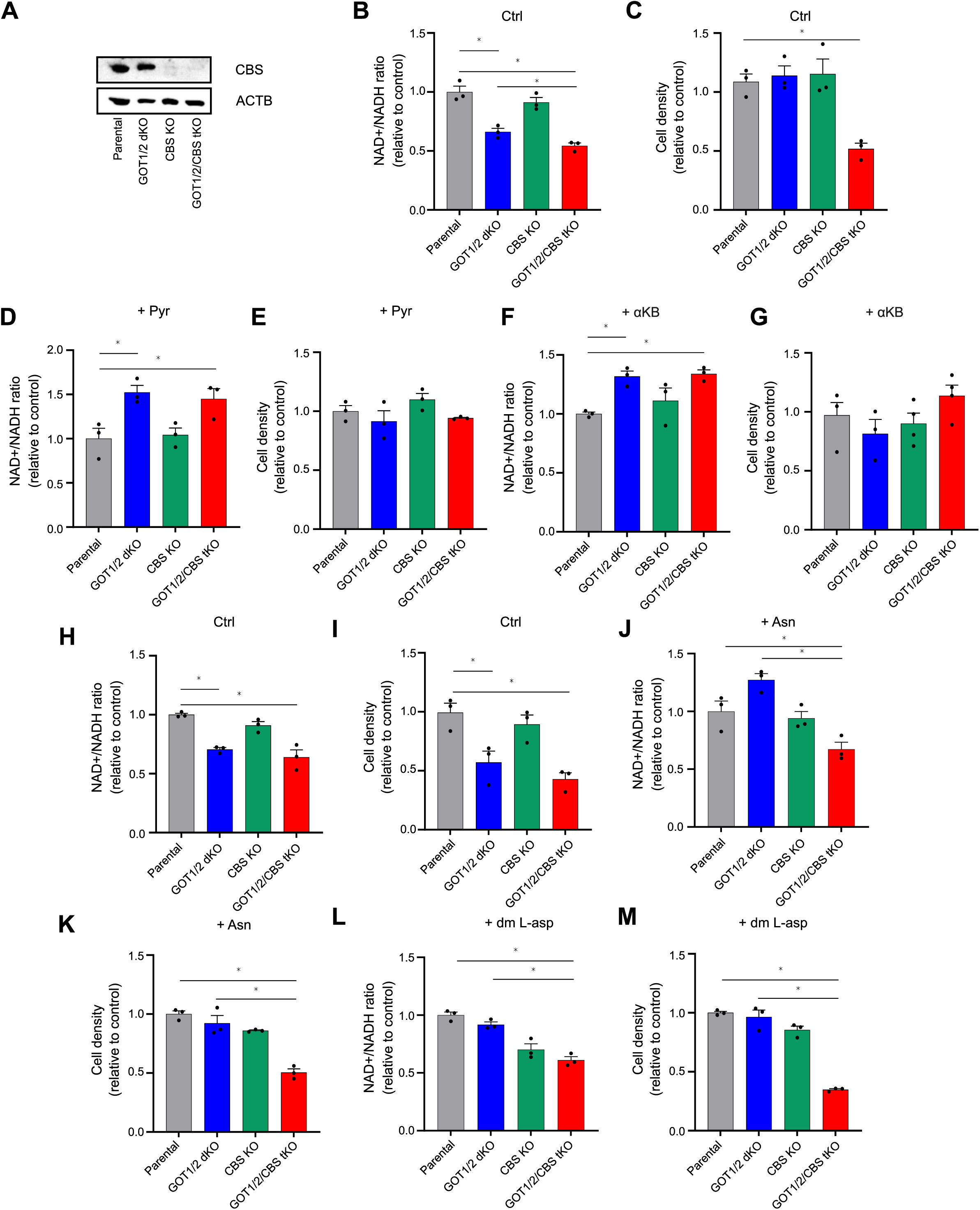
transsulfuration supports GOT1/2 dKO cells and enables rescue by asparagine. **A**) Confirmation of target protein ablation in GOT1/2/CBS tKO and CBS KO determined by west-ern blot. **B**) NAD+/NADH ratio in parental, GOT1/2 dKO, CBS KO, GOT1/2/CBS tKO cells grown in RPMI medium, measured by bioluminescence assay (mean ± SEM, n = 3, ∗p < 0.05, unpaired two-tailed t test). **C**) Proliferation in parental, GOT1/2 dKO, CBS KO, GOT1/2/CBS tKO cells grown in RPMI medium, measured by crystal violet assay after 72 h (mean ± SEM, n = 3, ∗p < 0.05, unpaired two-tailed t test). **D**) NAD+/NADH ratio in parental, GOT1/2 dKO, CBS KO, GOT1/2/CBS tKO cells grown in RPMI medium, in the presence of pyruvate (1 mM) measured by bioluminescence assay (mean ± SEM, n = 3, ∗p < 0.05, unpaired two-tailed t test). **E**) Proliferation in parental, GOT1/2 dKO, CBS KO, GOT1/2/CBS tKO cells grown in RPMI medium, in the presence of pyruvate (1 mM) measured by crystal violet assay after 72 h (mean ± SEM, n = 3, ∗p < 0.05, unpaired two-tailed t test). **F**) NAD+/NADH ratio in parental, GOT1/2 dKO, CBS KO, GOT1/2/CBS tKO cells grown in RPMI medium, in the presence of αKB (1 mM) measured by bioluminescence assay (mean ± SEM, n = 3, ∗p < 0.05, unpaired two-tailed t test). **G**) Proliferation in parental, GOT1/2 dKO, CBS KO, GOT1/2/CBS tKO cells grown in RPMI medi-um, in the presence of αKB (1 mM) measured by crystal violet assay after 72 h (mean ± SEM, n = 3, ∗p < 0.05, unpaired two-tailed t test). **H**) NAD+/NADH ratio in parental, GOT1/2 dKO, CBS KO, GOT1/2/CBS tKO cells grown in DMEM medium, measured by bioluminescence assay (mean ± SEM, n = 3, ∗p < 0.05, unpaired two-tailed t test). **I**) Proliferation in parental, GOT1/2 dKO, CBS KO, GOT1/2/CBS tKO cells grown in DMEM medi-um, measured by crystal violet assay after 72 h (mean ± SEM, n = 3, ∗p < 0.05, unpaired two-tailed t test). **J**) NAD+/NADH ratio in parental, GOT1/2 dKO, CBS KO, GOT1/2/CBS tKO cells in DMEM medium, in the presence of asparagine (340 µM) measured by bioluminescence assay (mean ± SEM, n = 3, ∗p < 0.05, unpaired two-tailed t test). **K**) Proliferation in parental, GOT1/2 dKO, CBS KO, GOT1/2/CBS tKO cells in DMEM medium, in the presence of asparagine (340 µM) measured by crystal violet assay after 72 h (mean ± SEM, n = 3, ∗p < 0.05, unpaired two-tailed t test). **L**) NAD+/NADH ratio in parental, GOT1/2 dKO, CBS KO, GOT1/2/CBS tKO cells in DMEM medium, in the presence of dimethyl L-aspartate (2 mM) measured by bioluminescence assay (mean ± SEM, n = 3, ∗p < 0.05, unpaired two-tailed t test).**M**) Proliferation in parental, GOT1/2 dKO, CBS KO, GOT1/2/CBS tKO cells in DMEM medium, in the presence of dimethyl L-aspartate (2 mM) measured by crystal violet assay after 72 h (mean ± SEM, n = 3, ∗p < 0.05, unpaired two-tailed t test).

In DMEM medium, which does not contain asparagine, both GOT1/2/CBS tKO and GOT1/2 dKO cells had a reduced NAD^+^/NADH ratio and proliferation rate when compared to parental or CBS KO cells (Fig. 6H, I), suggesting that transsulfuration cannot restore cellular redox or proliferation in the absence of asparagine. Consistently, asparagine and dimethyl aspartate rescued NAD^+^/NADH ratio and proliferation of GOT1/2 dKO cells, but they could not compensate GOT1/2/CBS tKO cells defects (Fig.6J-M). Together, these data support a model in which aspara-gine engages transsulfuration to contribute to cytosolic redox restoration and pyruvate-independent proliferation.

### TRANSSULFURATION SUPPORTS GOT1/2 DEFICIENT TUMOR GROWTH AND REPRESENTS A TARGETABLE VULNERABILITY IN CANCER

In contrast to defined in vitro conditions, the tumor microenvironment provides multiple redox-modifying substrates. For example, pyruvate and α-ketobutyrate, which normalize cytosolic redox in the absence of GOT1 and GOT2, were depleted (we interpret this as consumed) from the environment of GOT1/2 dKO tumors (c.f. Fig. 1I). Because transsulfuration enables proliferation in the absence of pyruvate and α-ketobutyrate in vitro, we tested whether it provides a similar tumor growth advantage in vivo. Cysteine, a product of transsulfuration, trended higher in GOT1/2 dKO tumors, indicating that activation of transsulfuration in the absence of GOT1/2 may be maintained in vivo (Fig 7A). We next treated mice bearing parental and GOT1/2 dKO melanoma tumors with the transsulfuration inhibitor AOA. Consistent with in vitro data, AOA administration reduced the growth of GOT1/2 dKO, but not parental tumors (Fig. 7B, C). Metabolic profiling revealed increased accumulation of cystathionine, a transsulfuration pathway intermediate, in the interstitial fluid of AOA-treated GOT1/2 dKO tumors compared to AOA-treated parental tumors (Fig. 7D). Similarly, genetic knockout of CBS impaired the growth of GOT1/2 deficient, but not parental tumors (Fig. 7E). The transsulfuration pathway thus promotes growth of GOT1/2 dKO tumors with compromised cytosolic redox state even when pyruvate and α-ketobutyrate are available.

**Figure 7:**
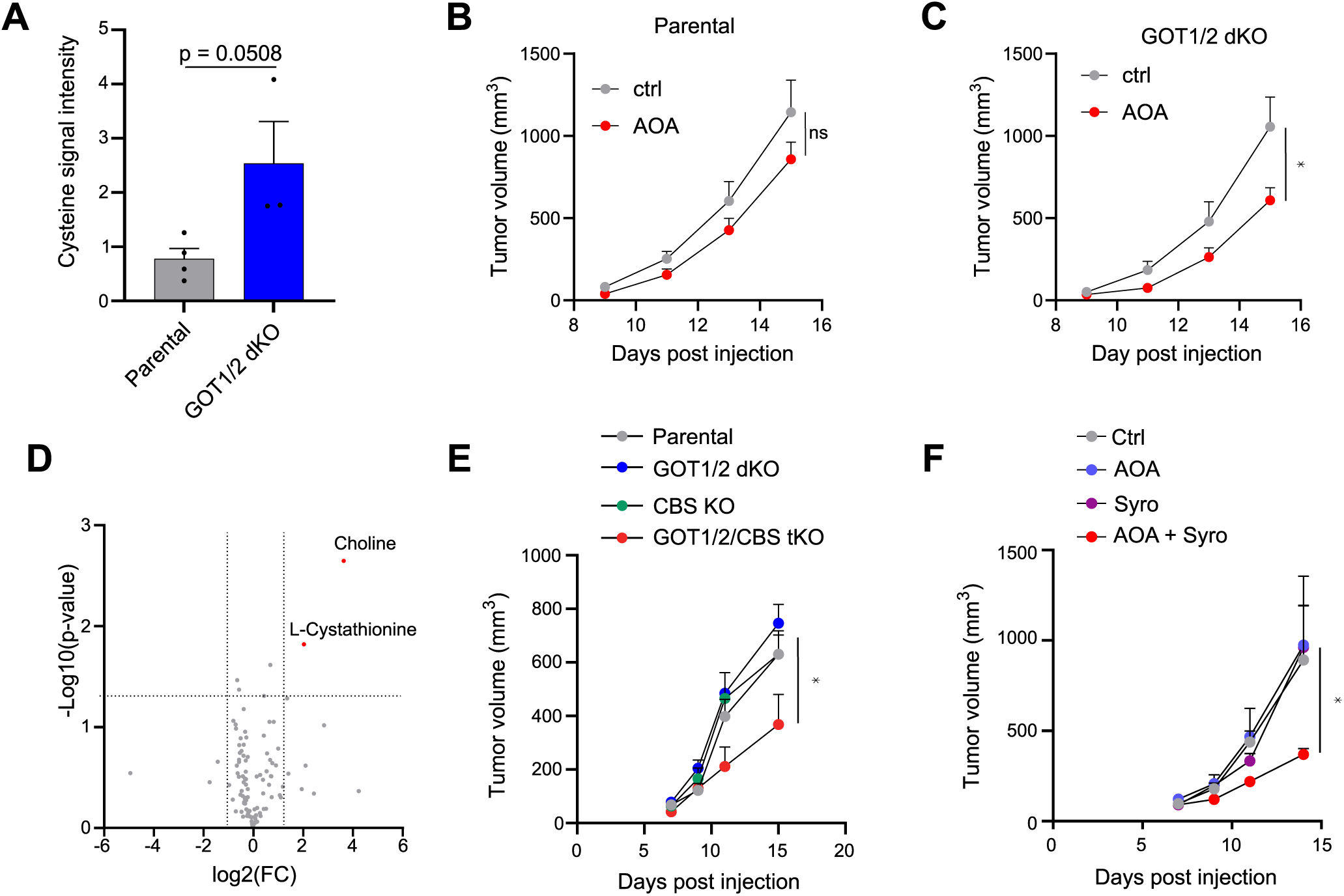
transsulfuration supports tumor growth. **A**) Intratumoral cysteine levels in parental and GOT1/2 dKO melanoma tumors, determined by MALDI imaging (mean ± SEM, n > 3, unpaired two-tailed t test). **B**) Tumor growth kinetics of mouse melanoma tumors generated by s.c. injection of parental cells, treated with 1.1 mg/kg aminooxyacetic acid (AOA) or with vehicle (PBS) every 24 h (mean ± SEM, n = 5, ∗p < 0.05, two-way ANOVA). **C**) Tumor growth kinetics of mouse melanoma tumors generated by s.c. injection of GOT1/2 dKO cells, treated with 1.1 mg/kg aminooxyacetic acid (AOA) or with vehicle (PBS) every 24 h (mean ± SEM, n = 5, ∗p < 0.05, two-way ANOVA). **D**) Quantification of metabolites in tumor interstitial fluid isolated from GOT1/2 dKO and paren-tal melanoma tumors treated with 1.1 mg/kg aminooxyacetic acid (AOA) measured by LC-MS. Blue indicates significantly decreased metabolites and red indicates significantly increased metabolites in GOT1/2 dKO tumors relative to parental B16 (n=3, significant hits log2FC > 1, p < 0.05). **E**) Tumor growth kinetics of mouse melanoma tumors generated by s.c. injection of parental, GOT1/2 dKO, CBS KO, GOT1/2/CBS tKO cells (mean ± SEM, n = 5, ∗p < 0.05, two-way ANOVA). **F**) Tumor growth kinetics of mouse melanoma tumors generated by s.c. injection of parental cells, treated with 1.1 mg/kg of aminooxyacetic acid (AOA), 5.5 mg/kg syrosingopine (SYR, a MCT1/4 inhibitor), their combination or with vehicle (corn oil) every 24 h (mean ± SEM, n = 5, ∗p < 0.05, two-way ANOVA).

Given the contribution of transsulfuration to cytosolic redox maintenance, we tested whether targeting transsulfuration-supported cytosolic redox could be exploited for cancer therapy. Effective modulation of the cytosolic redox state, without targeting transsulfuration, has not been achieved in vivo^20^. We treated parental melanoma tumors with AOA, which blocks both transsulfuration and the MAS, and syrosingopine, a pyruvate/α-ketobutyrate uptake blocker. While single agents had no effect, the combination treatment impaired tumor growth (Fig. 7F). Accordingly, targeting of the cytosolic redox state by simultaneously interfering with external (pyruvate/α-ketobutyrate uptake) and internal (MAS and transsulfuration) mechanisms of redox homeostasis represents an actionable strategy for cancer therapy.

## Discussion

This study identifies the cytosolic redox state, rather than aspartate availability, as the primary metabolic bottleneck in tumors lacking both GOT1 and GOT2. These two aspartate aminotransferases are core components of the malate-aspartate shuttle, which produces aspartate and maintains cytosolic NAD⁺/NADH balance. By disrupting both enzymes simultaneously, we were able to uncouple aspartate biosynthesis from redox maintenance. Unexpectedly, aspartate availability is not limiting. Instead, tumor growth in this context is supported by restoration of cytosolic redox state. This restoration is consistent with asparagine enabling serine/methionine-dependent transsulfuration to generate α-ketobutyrate, whose conversion to α-hydroxybutyrate can recover cytosolic NAD^+^ and sustain proliferation. Furthermore, the adaptation described here establishes a previously unrecognized two-way relationship between aspar-tate-asparagine metabolism and cytosolic redox control and shows that asparagine regulates the redox state in a MAS-independent manner. This identifies a new role for extracellular asparagine, which complements its previously described proteinogenic, anti-apoptotic and signaling functions^16,31,32^, and its contribution to preserving the intracellular aspartate pool^33^, in support of tumor growth.

In some contexts, other biosynthetic pathways could be impacted by redox defects more than aspartate synthesis^9,34,35^, underscoring the prime importance of redox balance as a limitation for proliferating cells that likely varies across cancer contexts. Prior work has placed aspartate shortage downstream of cytosolic redox control^6,7^, largely because restoring the NAD⁺/NADH ratio via exogenous electron acceptors rescues aspartate synthesis and proliferation. Our results extend these findings by showing that cancer cells can engage an alternative redox-balancing pathway through asparagine-induced transsulfuration, bypassing the MAS choke point. This bypass requires asparagine-enabled serine import, linking earlier observations on amino acid exchange^25^ directly to redox homeostasis. Aspartate contributes to this circuit only as the precursor to asparagine, in a MAS-independent manner. A recent report shows that aspartate also supports cytosolic redox directly, by driving NADH consumption through the GOT1–MDH1 arm of the MAS^22^. Together these findings define two routes by which aspartate-asparagine metabolism feeds cytosolic NAD⁺ regeneration, one MAS-dependent and one MAS-independent. In consequence, aspartate shortage in redox-compromised contexts could initiate a self-reinforcing decline in the cytosolic NAD⁺/NADH ratio, broken once asparagine becomes available to engage transsulfuration.

Our in vivo tumor data argue that the transsulfuration-mediated bypass operates even when exogenous electron acceptors are present in the tumor environment, suggesting this is a major compensatory route. Given that transsulfuration is traditionally studied in the context of cysteine and glutathione synthesis or hydrogen sulfide generation^24,36^, its role in producing α-ketobutyrate as a redox-active metabolite warrants broader consideration, beyond cancer. Upregulation of transsulfuration in models of partial mitochondrial insufficiency (e.g., mutator mouse)^27,37^ suggests that this adaptive circuit may be a general mechanism that cells use to maintain cytosolic redox rather than a GOT1/2 loss- or cancer-specific phenomenon.

Our metabolic CRISPR screen points to several components of mitochondrial oxidative phosphorylation as selectively essential in the absence of GOT1/2, suggesting that mitochondrial respiration remains necessary for asparagine-driven redox rescue by transsulfuration. Previous reports showed that asparagine supplementation does not normalize the NAD^+^/NADH ratio when respiration is inhibited^8,16^, and that cytosolic MAS enzyme defects (GOT1 and MDH1) are better tolerated than those in the mitochondria (GOT2, MDH2)^22^. Results presented here also indicate that severe complex I defects are harder to compensate in vivo than combined GOT1/2 deficiency. Together, this suggests that asparagine-dependent transsulfuration may not be able to rescue redox under complete respiratory insufficiency^6,7,16^. However, a recent report proposed that transsulfuration-derived α-ketobutyrate enables glucose oxidation and stimulates proliferation of cells with mitochondrial DNA mutations that do not fully suppress respiration (these cells did not feature a lower NAD^+^/NADH ratio or pyruvate auxotrophy, and asparagine was not present in the culture medium)^27^. Partial mitochondrial defects could thus still be compensated by transsulfuration, but the mechanism responsible might be different.

From a therapeutic perspective, our data suggest that direct pharmacological targeting of aspartate synthesis for cancer therapy may not be effective, as multiple routes can supply aspartate and sustain proliferation^17,20,38^. Targeting cytosolic redox directly may be more effective, as shown by the synergy between pyruvate uptake inhibitors and aminooxyacetic acid (AOA), which blocks GOT1, GOT2, and transsulfuration (as well as other pyridoxal phosphate-containing enzymes). However, novel inhibitors with improved specificity and safety profiles are needed. Furthermore, combining GOT1-specific inhibitors with transsulfuration blockade could offer a more selective cytosolic redox-targeted strategy, potentially avoiding the systemic toxicity of respiratory chain inhibition^39^ and minimizing the impact of losing mitochondrial ATP production in non-cancerous cells^40^. Beyond cancer, our results suggest that raising asparagine levels, possibly via diet, might enhance transsulfuration and improve outcomes in mitochondrial diseases with partial oxidative phosphorylation or MAS defects^37^.

Regardless, we show that asparagine-driven transsulfuration provides a robust bypass for cytosolic redox restoration in tumors, reframing the role of aspartate-asparagine metabolism in proliferation and revealing new avenues for metabolic intervention to exploit the tumor’s dependence on cytosolic redox balance.

## Limitations

Our data suggest an alternative aspartate synthase activity by an unknown enzyme(s). While this is of interest, we leave it up to future research to elucidate.

## Methods

### Cell Lines and Culture Conditions

B16-F10 mouse melanoma and 4T1 mouse mammary carcinoma were obtained from ATCC. HKP1 mouse lung adenocarcinoma cells were a kind gift of Prof. Vivek Mittal (Weill Cornell Medicine, USA). Parental cells were cultured in RPMI 1640 (Sigma-Aldrich) (B16-F10, 4T1 and HKP1). For routine culture, GOT1/2 dKO cells, CBS KO, and GOT1/2 CBS tKOs (see below) were maintained in a medium supplemented with 1mM sodium pyruvate (P5280, Sigma-Aldrich). ASNS KOs and GOT1/2 ASNS tKOs were maintained in medium supplemented with 1 mM sodium pyruvate (P5280, Sigma-Aldrich) and 340 µM of asparagine (A4284, Sigma-Aldrich). For nutrient dependency experiments, we mixed individual components to produce custom-made RPMI media lacking specific amino acids, as described in Table S6. Assay-specific media are detailed below. All cultures were regularly tested for Mycoplasma infection using the MycoAlert (Lonza) test.

#### Knockout generation

To generate GOT1/2 dKO in B16 and 4T1 cells, we used CRISPR/spCas12 (also known as CPF1) plasmid-based method. Arrays of 3 guide RNAs (gRNAs), selected by CRISPOR (https://crispor.gi.ucsc.edu/), were designed to remove/frameshift critical exons (exon 2 of GOT1 and exon 2 of GOT2). The sgRNA arrays (separate for GOT1 and GOT2, Table S7), were synthesized (Sigma-Aldrich), and cloned into the BbsI site of an AsCpf1-Venus-NLS crRNA entry plasmid (a kind gift of Dr. B. Schuster, Institute of Molecular Genetics of the Czech Academy of Sciences) using fast digest BpiI (FD1014, Thermo Fisher Scientific) and verified by sequencing. Cells were transfected using Lipofectamine 3000 (L300000, Invitrogen) according to the manufacturer’s protocol. After 2 days, Venus-positive cells were single cell sorted into 96-well plates and expanded in RPMI medium supplemented with 1mM sodium pyruvate and 204 μM uridine. Two rounds of this procedure were performed to obtain the GOT1/2 dKO. Selected clones were validated for gene deletion by PCR (for primers see Table S7) and western blotting.

To obtain GOT1/2 dKO in HKP1 cells, and ASNS and CBS KOs (and triple KOs) in B16 cells we used ribonucleoprotein (RNP)-based editing. Cells were electroporated with pre-assembled gRNA-Cas9 RNP complexes (Gene Knockout Kit v2, Synthego, see Table S7 for sequences) using the Neon™ Transfection System (Thermo Fisher Scientific) with 10 μL Neon™ tips. Electro-poration was performed at 1150 V, 30 ms pulse width, and 2 pulses for HKP1 cells and 1400 V, 20 ms pulse width, 2 pulses for B16 cells. Following electroporation, cells were pelleted in two halves. The first half was used to confirm gene editing efficacy in bulk (PCR and sequencing) while the other one was cryopreserved. When successful editing was confirmed, cryopreserved cells were recovered, single-cell sorted into 96-well plates and expanded as above. Target gene deletion in selected clones was confirmed by PCR, sequencing and western blotting.

### Stable expression of cytoplasmic and mitochondrial LbNOX

Stable expression of cytoplasmic and mitochondrially targeted Lactobacillus brevis NADH oxidase (LbNOX) was achieved using the pCDH lentiviral system. The pUC57 vectors containing cytoplasmic LbNOX and mitochondrial LbNOX (mtLbNOX) were a gift from Vamsi Mootha. The LbNOX and mtLbNOX coding sequences were PCR-amplified using iProof DNA Polymerase (BioRad) with primers introducing EcoRI and BamHI restriction sites and a C-terminal FLAG tag. PCR products were purified, digested with EcoRI/BamHI, and ligated into the pCDH-CMV-MCS-EF1-Puro vector using T4 DNA ligase (Thermo Fisher). Positive clones were verified by Sanger sequencing. Plasmids were transfected into B16 cells using Lipofectamine 3000 (Thermo Fisher), and stable pools were selected with 2 µg/mL puromycin and maintained in 0.5 µg/mL for rou-tine culture.

### Mouse Models of tumor growth

B16 and 4T1 cancer cells (2.5 × 10^5^) were resuspended in 100 µl PBS and injected subcutaneously into the right flank of syngeneic mice (males and females C57BL/6 for B16, females BALB/c for 4T1 cells, 7–12-week-old) using a 26G needle. Tumors were measured every 2–3 days using calipers and tumor volume was calculated as (length × width²)/2. For pharmacological treatments, animals with palpable tumors were randomized to receive daily intraperitoneal injection of AOA (1.1 mg/kg in PBS), syrosingopine (5.5 mg/kg in corn oil), or solvent control. Tumor growth and body weight were monitored until tumors reached the ethical limit (1000 mm^3^).

HKP1 cells expressing firefly luciferase were used to generate orthotopic tumors upon tail vein injection (1 × 10^5^). Bioluminescence imaging was performed after intraperitoneal injection of D-luciferin (15 mg/mL, 10 μL/g body weight, Revvity) followed by isoflurane anesthesia. Images were acquired using an Aura Lago X Spectral system (Spectral Instruments Imaging) with 300s exposure, binning = 4, f-stop = 1.2, field of view = 25. Low energy X-ray reference images were recorded. Data was analyzed using Aura software and reported as total emission (pho-tos/second).

Animals were obtained from established breeding colonies of the Czech Center of Phenogenomics (Institute of Molecular Genetics of the Czech Academy of Sciences) and were maintained under SPF conditions in individually ventilated cages with controlled temperature (22 ± 2°C) and humidity under a 12 h light/12 h dark cycle and with food and drink ad libitum. Animal health was monitored daily by experimenters, caretakers, and veterinarians. All animal experiments were approved by the Animal Ethics Committee of the Czech Academy of Sciences and were performed according to the Czech guidelines for the Care and Use of Animals in Re-search and Teaching.

### Cell proliferation assays

CRYSTAL VIOLET STAINING: Cells (2.5 x 10^3^) were seeded in 96-well plates in 100 μL of growth medium. After attachment, cells were washed with PBS, and the medium was replaced with experimental conditions using dialyzed FBS (S181D-500, Biowest). The plates were fixed at 72 h with 60 μL of 4% paraformaldehyde (9713.1000, VWR) per well for 30 min at room temperature (RT). After washing with PBS, cells were stained with 0.05% Crystal Violet (C0775, Sigma-Aldrich) for 11h, washed again, and the dye was solubilized in 1% SDS (L3771, Sigma-Aldrich). Absorbance was read at 5951nm (Infinite M200, Tecan). Conditions were tested in quadruplicates unless otherwise specified.

REAL-TIME CELL PROLIFERATION ASSAYS – CYTATION5: Cells (2 x 10^3^-2.5 x 10^3^ per well) were seeded in 96-well plates in 100 μL of standard growth medium. After attachment, cells were washed with PBS, and the medium was replaced with experimental conditions. Each condition was performed in triplicates unless otherwise specified. Growth was monitored over 72 h using a Bio-Tech Cytation5 imager with Gen5 software under 5% CO_2_ and atmospheric O_2_. INCUCYTE: Cells (2.5 x10^3^ cells/well) were seeded in 96-well plates and monitored every 41h for 721h using IncuCyte SX1 analyzer (Sartorius & Essen BioScience). Images were scanned with 10x objective every 4 h and analyzed with Incucyte Analysis Software.

Cells were routinely maintained in RPMI 1640 supplemented with 10% fetal bovine serum (FBS) and 1 mM sodium pyruvate. Prior to experiments, cultures were switched to the indicated experimental media. Where specified, RPMI 1640 supplemented with 10% dFBS was used as the basal medium. For DMEM-based conditions, DMEM (D5030; Sigma-Aldrich) was supplemented with 2 g/L sodium bicarbonate, 11 mM glucose, 2 mM L-glutamine, and 10% dFBS. Custom RPMI formulations were prepared by reconstituting all components to standard RPMI concentrations, with selective omission or adjustment of individual components as indicated for specific exper-imental conditions.

### Metabolite Profiling in cell culture

#### Steady state metabolite profiling by GC-MS

Cells were plated in 6-well plates, including cell-free wells for background correction. Cell number normalization was performed in parallel wells using standard cell counting and BSA protein quantification. Cells were cultured for 24 h in medium containing dialyzed FBS, washed three times with PBS, and subsequently maintained in culture medium formulated with the indicated experimental conditions. At the collection point wells were washed 2–4 times with cold 0.9%

NaCl, placed on ice, and quenched with 500 μL of ice-cold extraction solvent per well (80% MeOH containing internal standard d6-glutaric acid). Cells were scraped quickly and transferred to pre-chilled Eppendorf tubes on wet ice, vortexed for 2 min on 4°C and stored overnight at −80°C. Samples were centrifuged at 14,0001×1g for 15 min at 4°C, and 300–400 μL of the polar supernatant was transferred into pre-chilled, labeled tubes. The polar extracts were evaporated to dryness using a SpeedVac at 37°C for ∼2 h. For medium samples, 100 μL of thawed culture supernatant was mixed with 400 μL of extraction solvent (100% MeOH) and vortexed briefly. Samples were stored overnight at −80°C, centrifuged (14,0001×1g, 151min, 4°C), and the polar supernatant was collected and evaporated as described. <u>Sample derivatization:</u> Dried metabolite extracts were derivatized by adding 16 μL methoxamine reagent (TS-45950, Thermo Fisher Scientific) and incubated for 1 h at 37°C. Subsequently, 20 μL N-tert-butyldimethylsilyl-N-methyltrifluoroacetamide containing 1% tert-butyldimethylchlorosilane (375934, Sigma-Aldrich) was added, and samples were incubated for an additional 2 h at 60°C. After derivatization, samples were centrifuged at maximum speed for 10 min, and 20 μL of the supernatant was used for GC–MS analysis. <u>Data processing</u>: Derivatized samples were analyzed using a DB-35MS column (30 m × 0.25 mm i.d. × 0.25 μm; Agilent J&W Scientific) installed in an Agilent 7890 gas chromatograph coupled to an Agilent 5975C mass spectrometer. Ion counts for each detected metabolite were normalized to the internal standard norvaline and to cell number, which was determined using a Cellometer. This method was used in Fig. 2 B,C,G and H.

### METABOLITE PROFILING BY LC-MS

For in vitro intracellular metabolite labeling measurements, RPMI medium (without cystine, asparagine, serine and methionine) was supplemented with 10% dialyzed FBS and 100 µM U-^13^C5-Methionine- and 285 µM unlabeled serine, or with 285 µM U-^13^C3-Serine. B16 cells were seeded at a concentration of 7 x 10^5^ cells/mL, and a total of 5 x 10^6^ cells were used per replicate for each condition. The cells were supplemented or not with 340 μM asparagine and incubated with the tracers for 24 h. For the asparagine/serine exchange experiment, cells were preincubated with 340 μM asparagine in the absence of serine for 2 h, and then the medium was switched to asparagine free medium containing 285 µM U-^13^C3-Serine for 2 h. At endpoint, the medium was discarded and cells were rinsed for a few seconds with approx. 2 mL of ice-cold Blood Bank Saline. The plate was then laid on liquid nitrogen to quickly quench cellular metabolism and then transferred onto a mix of ice/dry ice (1:1). Intracellular metabolites were extracted with ice-cold 800 µL of MeOH/H2O (8:2, v:v) containing 5 mM N-ethylmaleimide, 0.1% formic acid, and 0.5 µg/mL glutarate and norvaline as internal standards. Next, the samples were vortexed at 4°C for 1.5 h and then centrifuged at maximum speed (16000 x g) at 4°C for 10 min. The supernatant containing the extracted metabolites was split equally in half (for the different MS methods) and dried overnight under vacuum at 4°C. Subsequently, samples were resuspended in 50 µL 80% methanol containing 0.1% formic acid, vortexed for 10 min at 4°C, and transferred into glass chromatography vials for mass spectrometry analysis.

For the LC-MS analysis, a 1290 Infinity II liquid chromatography (Agilent Technologies) with a thermal autosampler set at 4°C, coupled to a Q-TOF 6546 mass spectrometer (Agilent Technologies) was used. 5 µL sample were injected into an InfinityLab Poroshell 120 HILIC-Z, 2.1 x 150 mm, 2.7 µm (Agilent Technologies). Polar metabolites were analyzed in negative ionization mode, where the separation of metabolites was achieved using a gradient for 32 min at 50°C with a flow rate of 0.25 mL/min (solvent A: 10 mM ammonium acetate (pH=9) in water + 2.5 µM InfinityLab Deactivator Additive (Agilent); solvent B: 10 mM ammonium acetate (pH 9) in 85% acetonitrile + 2.5 µM InfinityLab Deactivator Additive (Agilent)); 0 min: 4% A, 2 min: 4% A, 5.5 min: 12% A, 8.5 min: 12% A, 9 min: 14% A, 14 min: 14% A, 19 min: 18% A, 25 min: 35% A, 27 min: 35% A, 28 min: 4% A, 32 min: 4% A). The MS operated in negative mode (m/z range: 50–1,200) using a sheath gas temperature of 350°C (12 L/min) and a gas temperature at 225°C (13 L/min). The nebulizer was set at 35 psi, the fragmentor at 125V, and the capillary at 3500 V. The analysis of metabolites of the transsulfuration pathway was performed in positive ionization mode, where the separation of metabolites was achieved at 25 °C with a flow rate of 0.25 mL/min. A gradient was applied for 30 min (solvent A: 10 mM ammonium formate in water (pH=3.8 adjusted with formic acid); solvent B: acetonitrile) to separate the targeted metabolites (0 min: 90% B, 2 min: 90% B, 12 min: 40% B, 15 min: 40% B; 25 min: 90% B; 30 min: 90% B). The mass spectrometer was operated in positive full scan mode (m/z range: 50–1,200) using a sheath gas temperature of 225 °C (10 L/min) and a gas temperature at 225 °C (10 L/min). The nebulizer was set at 40 psi, the fragmentor at 125 V and the capillary at 3,000 V. Data was analyzed using the Masshunter Profinder software (Agilent Technologies, v.10.0.2. build 10.0.2.162) and normalized by the internal standard and protein levels. Retention times for each metabolite were confirmed by single standards, which were prepared and analyzed in parallel to the biological samples. This method was used for Figures 5D (U-^13^C3 Serine tracing) and 5G (absolute values, contains U-^13^C5 methionine tracer).

Tracing into aspartate and aspartate uptake:

<u>Sample preparation:</u> Samples for 13C tracing were prepared following the steady-state sample collection protocol described above. Uniformly 13C-labeled tracers were used at RPMI concentrations in the absence of corresponding 12C substrates in DMEM/10% dialyzed FBS. Cells were then cultured under the specified experimental conditions. S<u>ample derivatization</u>: Dried metabolite extracts were derivatized using a two-step procedure. First, 20 μL of 2% (w/v) methoxyamine hydrochloride (89803, Sigma-Aldrich) in pyridine was added to the dried extracts and incubated at 60°C for 1 h. Samples were briefly centrifuged (10 s) to collect the solution at the bottom of the tube, followed by the addition of 30 μL of MTBSTFA containing 1% TBDMCS (Thermo Fisher). Samples were incubated for a further 1 h at 60°C, centrifuged at 3000 × g for 15 min, and the supernatant was transferred into amber GC–MS vials for analysis. <u>Analysis</u>: GC–MS analysis was performed using an Agilent 7890B GC system coupled to a 5977A MSD with a 30 m RXI-5MS capillary column. A 1 μL sample was injected in splitless mode with helium carrier gas at 1 mL/min and an inlet temperature of 270°C. The oven program consisted of an initial hold at 100°C for 1 min, a ramp to 160°C at 10°C/min, a ramp to 200°C at 5°C/min, and a final ramp to 330°C at 10°C/min with a 4 min hold. The transfer line was held at 270°C. Ionization was performed by electron impact (70 eV) with a source temperature of 230°C, and data were collected in scan mode.

<u>Data processing</u>: Spectra were processed in MATLAB (2024a) using the METRAN script. Mass isotopomer distributions (MIDs) were corrected for natural isotope abundance. Metabolite levels were normalized to the internal standard D6-glutaric acid and further corrected to protein abundance or cell counts. This method was used in Figure S2E, M and N.

### Metabolite profiling in tumor interstitial fluid and plasma

Tumors were excised, and the interstitial fluid was collected by centrifugation on nylon mesh filters as described^41^. In parallel, cardiac blood was collected for plasma isolation^41^. 5 μL of each sample or external chemical standard pool (ranging from ∼51mM to ∼11μM) was mixed with 45 μL of extraction solvent composed of acetonitrile:methanol:formic acid (75:25:0.1, v/v/v) which included isotopically labeled internal standards (see Materials section for details). The mixture was vortexed for 151min at 41°C to facilitate the extraction of polar metabolites. Insoluble material was pelleted by centrifugation at 16,0001×1g for 101min at 41°C. 201μL of super-natant containing the soluble polar metabolite fraction was used for LC-MS analysis. Instrumentation setup and measurement conditions were performed as previously described^42^.

Metabolite identification was performed using XCalibur 2.2 software (Thermo Fisher Scientific, Waltham, MA). Identification criteria included a mass accuracy threshold of 51ppm and a retention time window of 0.5 min. Assignment of metabolites was based on external standard pools, which were used to match observed peaks to known m/z and retention times, and to determine the limit of detection for each compound. Limits of detection ranged from 1001nM to 31μM (see Table S8 for retention time and m/z values for all analyzed metabolites). Quantification of metabolites was performed using two parallel approaches: (1) stable isotope dilution, and (2) external calibration. For metabolites with available isotopically labeled internal standards, quantification was performed by stable isotope dilution. First, peak areas of the internal standards were compared to those from serial dilutions of external standard pools to determine the absolute concentration of each internal standard in the extraction mix. Subsequently, the peak area of each unlabeled metabolite in the samples was compared to that of its corresponding quantified internal standard to determine metabolite concentration. Using this method, 63 metabolites were quantified. External calibration was used for metabolites lacking specific internal standards. Peak areas of these analytes were normalized to a labeled amino acid internal standard that eluted at a similar retention time, to correct for sample-to-sample variability in recovery. Normalized peak areas were then used to construct standard curves from the external standard pool dilutions. These curves, which showed linear relationships (typically with r² ≥ 0.95), were used to convert sample peak areas into absolute concentrations. Metabolites that did not meet these criteria were excluded from further analysis. Using this external calibration approach, an additional 44 metabolites were quantified (Table S8).

### Metabolite profiling in situ

<u>Tissue Collection:</u> Mice were sacrificed by CO_2_. Tumors were washed serially in pre-warmed 2 and 10% (w/w) gelatin solutions, respectively. The samples were then transferred to cryomolds with fresh 10% gelatin solution and flash-frozen in the 2-methylbutane tempered in liquid nitrogen and subsequently stored at - 80 °C prior to cryosectioning.

<u>Cryosectioning</u> Frozen tissue blocks were cooled to -12 °C in the CM1950 cryostat (Leica Biosystems) and cut in random order to 10 µm thick sections. Cryosections were mounted onto warm ITO glass slides (Ossila, UK) coated with 1:1 mixture (v/v) of poly-L-lysine (Sigma) and 0.1% of Nonidet P-40 Substitute (Sigma). Freshly attached cut on glass was left dried on hot plate at 40°C for 15 sec (LABOTECT Hot Plate 100, Labotect). and dried in a desiccator for 15 min, packed in plastic slide mailers, and vacuum-sealed for storage at -80 °C until MALDI imaging.

<u>Tissue slide preparation for MALDI imaging:</u> Tissue sections were removed from the freezer and adjusted to room temperature for 30 min and placed in a desiccator for 15 mins to dry. For measurements using the MALDI-Time-of-Flight instrument, 9-aminoacridine (9AA; 92817, Sig-ma-Aldrich) at 10 mg/ml in 70% ethanol (v/v) was sprayed onto tissue sections in 12 matrix layers using the following parameters: spray nozzle temperature, 50 °C; flow rate, 35 µl/min; horizontal nozzle movement speed, 1000 mm/min; nitrogen flow rate, 2 l/min; gas pressure, 10 psi; distance between each pass, 2 mm; nozzle distance, 42 mm.

<u>MALDI imaging measurement:</u> MALDI-TOF spectra were acquired using the rapifleX MALDI-TOF/TOF mass spectrometer (Bruker Daltonics) operated by the fleXcontrol v4.2 software (Bruker Daltonics) in the reflector negative-ion mode with the 355 nm SmartBeam™ 3D laser (spatial resolution of 50 × 50 µm, m/z range of 20–1000) at the constant laser fluence of 84% and laser frequency of 5 kHz. 200 shots were summed up from each position. The instruments were set up accordingly: ion source 1, 20.105 kV; PIE, 2.663 kV; lens, 11.336 kV; reflector 1, 20.804 kV; reflector 2, 1.034 kV; reflector 3, 8.575 kV. The pulsed ion extraction time was set to 90 ns, detector gain was 2302 V. Data were acquired with a digitizer speed at 2.5 GS/s. Calibration was done externally using red phosphorus to achieve precision up to 2 ppm ^43^. Spatial navigation for imaging data acquisition was done in the flexImaging v5.0 software (Bruker Daltonics).

<u>MALDI imaging data processing:</u> Raw data files generated by the TOF and FTICR instruments were imported as full spectra into SCiLS Pro software v2021c (Bruker Daltonics). Spectra from TOF measurements were normalized by the total ion current method (TIC Intensity at peak maximum was reported in both cases using the rainbow color map. Images were displayed with linear interpolation, and the image denoising level was set to “weak”. The hotspot removal level was set to the 99% quantile. For better image contrasting, 100% color intensity of the color map was used. Averaged spectra were exported as an ASCII file and imported into the mMass software v 5.5.0 ^44^. Spectra acquired by MALDITOF were internally re-calibrated utilizing known compounds, reaching mass accuracy of 5 ppm from www.hmdb.ca (v4) ^45^, and only features annotated by the provider as “quantified” and “detected” were considered for putative annotations. MALDI-TOF spectra were annotated within ± 10 ppm interval. Only [M-H]- ions were con-sidered.

<u>H&E staining:</u> After MALDI imaging, 9-aminoacridine layer was removed from glass and tissue in Coplin’s jar filled with 70% ethanol. After 5 min, sample was washed with distilled water and proceed to hematoxylin-eosin staining, dehydrated by series of ethanol, isopropanol and xylene, and mounted in permanent medium (Pertex, Bamed) at Leica ST5020 automated staining instrument in combination with the Leica CV5030 coverslipper (Leica Biosystems). Tissue was then scanned with Axio Scan.Z1 (Zeiss) operated by Zen 2 software (blue edition; Zeiss) and exported as PNG file.

### Western blotting

Cells were lysed in RIPA buffer (20 mM Tris-HCl (pH 7.5), 1501mM NaCl, 11mM EDTA, 11mM EGTA, 1% NP-40, 0.5% sodium deoxycholate, and 0.1% SDS), supplemented with protease and phosphatase inhibitor cocktails (1% v/v each). Lysates were cleared by centrifugation (≥12,0001×1g, 10–151min, 41°C), protein concentration measured by BCA assay, and 30 μg of protein per lane was separated by SDS-PAGE and transferred to nitrocellulose membranes. Membranes were blocked, incubated with primary antibodies of interest (Resources Table), and detected by HRP-conjugated secondary antibodies using chemiluminescence.

### Fluorescence lifetime imaging of NAD(P)H

B16 cells were cultured in glass-bottom microscopy plates in DMEM medium (10% dFBS) and pre-treated with 11mM pyruvate for 4 h. Live cell imaging for combined NAD(P)H autofluorescence/FLIM was done using a Zeiss LSM880 NLO inverted microscope equipped with a Chameleon Ultra II multiphoton laser and Compact OPO (Coherent) and a non-descanned BiG-2 GaAsP detector. Imaging was performed with a 63×/1.4 NA Plan-Apochromat oil immersion objective. Excitation was at 7401nm (81mW at the source, 61mW during scanning) and emission was collected between 390–480 nm.

Image segmentation was performed to indirectly identify cytosol, nuclei, mitochondria, and melanin granules for subsequent lifetime analysis. First, composite RGB images encoding mean arrival time and single-channel intensity images were generated from raw TTTR data. The subsequent segmentation pipeline was a two-stage process involving initial object identification using machine learning in FIJI, followed by refined segmentation with fine-tuned Cellpose models and mask processing in a custom Python script. Melanin and mitochondria were segmented using the Trainable Weka Segmentation plugin (v3.3.2) for FIJI/ImageJ. Nuclei and cells were segmented with fine-tuned Cellpose (v3) models.

For melanin granules, the RGB lifetime image was used as input for a pre-trained classifi-er Melanin structures were segmented using a FastRandomForest classifier (200 trees, 2 features per split) implemented in WEKA. Features included Gaussian, Difference of Gaussians, Sobel, Brightness, and Saturation, with sigma values ranging from 1 to 16. The resulting probability map was binarized (threshold = 0.5), and morphological dilatation (2 pixels) was applied to create the final melanin mask. For mitochondria, the intensity image was first smoothed with a mean filter (radius = 2.0 pixels) and then processed with a separate pre-trained Weka classifier (200 trees, 2 features per split). Feature set included Gaussian, Difference of Gaussians, Sobel, Hessian, and Membrane projections, with sigma values ranging from 1 to 16. The resulting probability map was similarly binarized. To ensure exclusivity, the previously generated melanin mask was subtracted from the mitochondria mask.

Cells were segmented from the intensity images using a fine-tuned Cellpose model (Ini-tialized with the cyto2 model. Training parameters: learning rate 0.1, weight decay 0.0001, and 100 epochs). To accommodate cell size variability, the model was run three times with estimated object diameters of 150, 200, and 250 pixels, all using a flow threshold of 0.7. The resulting masks from all three runs were merged to form a final cell mask. To segment nuclei, a processed nuclear-enhanced image was first generated. The red channel of the RGB image, corresponding to longer lifetimes, was isolated and smoothed with a Gaussian blur (sigma = 2.0 pixels). The intensity image was inverted, and the result was divided by the blurred red channel image, producing an image where nuclear regions, characterized by low intensity and long lifetimes, were highlighted. Images were additionally masked with the corresponding final cell mask to limit the search space and a fine-tuned Cellpose model was then applied to these images to segment nuclei. This model was also run three times with varying parameters to capture a range of nuclear sizes: diameter = 55 pixels (flow threshold = 0.9), diameter = 125 pixels (flow threshold = 0.7), and diameter = 155 pixels (flow threshold = 0.7). The three resulting nuclear masks were merged. Final mask processing was performed to ensure that each pixel was assigned to a single compartment. The merged nuclear mask was refined by filling any internal holes, and then both mitochondria and melanin masks were subtracted from it. The cytoplasm was defined as the area within the final cell mask after subtracting the refined nuclear, mitochondria, and melanin masks. This cytoplasmic mask was further refined by a morphological binary erosion (1 pixel). Mitochondria were masked with the final cell masks to remove any segmented objects located outside of cell boundaries.

### NAD+/NADH MEASUREMENT

Intracellular NAD^⁺^ and NADH levels were quantified using the NAD/NADH-Glo Assay (G9072, Promega). Briefly, cells were seeded in 96-well plates at an initial density of 10,000 cells/well and allowed to adhere overnight under standard culture conditions. The following day, cells were washed once with 1× PBS and incubated with 100 µL of treatment medium for the indicated durations before extraction. For extraction, cells were washed once with ice-cold 1× PBS and lysed in 100 µL ice-cold 1% dodecyltrimethylammonium bromide (DTAB) in 0.2 N NaOH, diluted 1:1 with 1× PBS. Lysates were immediately flash-frozen in liquid nitrogen and stored at –80 °C until analysis. To measure NADH, 10 µL of each extract was transferred to 96 wells, mixed with 10 µL DTAB, and incubated at 75 °C for 30 min to selectively degrade NAD⁺ under basic conditions. To measure NAD^⁺^, 10 µL of extract was mixed with 30 µL DTAB and 20 µL 0.4 N HCl, followed by incubation at 60 °C for 15 min to degrade NADH under acidic conditions. After incubation, samples were equilibrated to room temperature and neutralized by adding 20 µL 0.25 M Tris in 0.2 N HCl (Penta) (for NADH-treated samples) or 20 µL 0.5 M Tris base (for NAD^⁺^-treated samples). Subsequent luminescence-based quantification was performed according to the manufacturer’s instructions using a Tecan Infinite M200 Pro luminometer.

### CRISPR SCREEN

AMPLIFICATION OF MOUSE CRISPR METABOLIC GENE KNOCKOUT LIBRARY: Electrocompetent Endura E. coli (60242-1, Lucigen) were transformed with 100 ng of pooled CRISPR sgRNA of mouse metabolic library (Addgene #160129, a gift of K. Birsoy) and electroporated at 1.81kV in 11mm cu-vettes (MicroPulser Electroporator, Biorad). Each reaction was immediately resuspended in recovery medium (80026-1, Lucigen), combined to a final volume of 4 mL, and incubated at 371°C for 11h with shaking. To quantify transformation efficacy, 10 μL of the transformation mixture was serially diluted (1:10) and plated on 10 cm LB-agar plates containing carbenicillin (205805, Millipore). The remaining 4 mL of culture were plated into two 245 mm square LB-carbenicillin plates and incubated overnight at 37 °C. Colonies were harvested in 20 mL of cold LB per plate, pooled, and centrifuged at 60001×1g for 151min. Plasmid DNA was extracted from the bacterial pellet using a maxi-prep kit (Qiagen) and quantified by Nanodrop.

LENTIVIRAL LIBRARY PACKAGING: 293FT cells were maintained in DMEM high glucose (D6429, Sigma-Aldrich) supplemented with 10% FBS and penicillin/streptomycin under standard conditions (37 °C, 5% CO₂). At confluency, 4 × 10⁶ cells were seeded into a 100-mm tissue-culture dish. For transfection, plasmid DNA was prepared in 500 µL Opti-MEM by mixing 16 µL Lipofectamine P3000 reagent (Invitrogen, L3000008) with 3.9 µg amplified library plasmid, 2.6 µg psPAX2, and 1.3 µg pMD2.G (total 7.8 µg DNA). In a separate tube, 12 µL Lipofectamine 3000 (Invitrogen, L3000008) was diluted in 500 µL Opti-MEM. Both mixtures were briefly vortexed and incubated at room temperature (RT) for 5 min. The DNA/P3000 mixture was then combined with the Lipofectamine mixture, vortexed briefly, and incubated a further 15 min at RT to allow complex formation.

Cells were gently resuspended in 6 mL pre-warmed Opti-MEM, and the transfection complexes were added dropwise. Cultures were returned to the incubator for 6 h, after which the medium was replaced with 14 mL fresh complete DMEM. Supernatants were harvested 48 h post-transfection, clarified by centrifugation at 800 rpm for 5 min at 4 °C, and passed through a 0.45 µm filter. PEG-it™ Virus Precipitation Solution (LV810A-1, System Biosciences) was added to the clarified supernatant at a 1:4 ratio (e.g., 3.5 mL PEG-it per 14 mL supernatant), mixed, and incubated overnight at 4 °C on a rotator. Virus was pelleted by centrifugation at 1,800 rpm for 40 min at 4 °C, resuspended in PBS at a 1:150 pellet:PBS ratio, aliquoted, and stored at −80 °C.

LENTIVIRAL TRANSDUCTION: B16 parental and GOT1/2 dKO knockout cells were thawed from liquid nitrogen stocks, recovered in culture for 1–2 days, trypsinized and seeded at 2.51×110⁵1cells per well in 2 mL of complete RPMI medium (10% FBS, P/S, 1 mM sodium pyruvate and 204 μM uridine) in 6-well plates (8 wells per cell type) to achieve 100x coverage per sgRNA. The following day, medium was replaced with 1.5 mL of fresh complete RPMI containing 8 μg/mL polybrene (TR-1003-G, Millipore). The cells were transduced at MOI 0.3-0.5 with lentiviral particles carrying the CRISPR library, and plates were centrifuged (8001×1g for 901min, 321°C), followed by a 6 h incubation at 37 °C with 5% CO₂. Next, medium was replaced with 2 mL of fresh complete RPMI without polybrene, and cells were incubated for an additional 24 h. Cells were trypsinized, split into T25 flasks, and selected with 30 μg/mL puromycin (ant-pr-1, Invivogen) for 31days at 371°C, 5% CO₂. Following selection, cells (7.5 x 10) were collected for DNA isolation to establish a baseline (Day 0).

LOSS OF FUNCTION SCREEN AND DATA ANALYSIS: Cells in the remaining plates were maintained for 14 doublings under the experimental condition (RPMI, 10% dialyzed FBS, P/S, ± 1 mM pyruvate). After that, 10 x 10^6^ cells per condition were collected for genomic DNA isolation (DNeasy Blood & Tissue Kit, Qiagen, 69506, Qiagen). 16 μg of isolated genomic DNA (8 parallel reactions, 2 µg gDNA/reaction) was used for sgRNAs amplification and index/Illumina adaptor insertion in a one-step PCR (Herculase II, Agilent, see Table S9 for primers and PCR conditions). The PCR product (± 300 bp band) was gel-purified (GeneJET PCR Purification Kit, K0701, Thermo Fisher Scientific) and its homogeneity was confirmed by the fragment analyzer (Fragment Analyzer 5200, Agilent). Barcoded products were pooled and used for sequencing at Illumina NextSeq™ 2000 using the NextSeq 1000/2000 P1 Reagent Kit.

DATA ANALYSIS: Sequencing quality from CRISPR screens performed in B16 parental and B16 GOT1/2 dKO cells was assessed using MultiQC v1.24 . Reads were processed with MAGeCK v0.5.9.5 using default parameters. Guide-level counts were generated separately for parental and dKO samples with the MAGeCK count module and normalized to a set of 50 intragenic control sgRNAs. Differential sgRNA abundance was estimated using the MAGeCK mle module with design matrices specifying day 0 as baseline versus experimental conditions. Quality control of MAGeCK outputs was performed with VISPR v0.4.14. Further statistical analyses were conducted in R v4.3.2. Genes with FDR < 0.25 were considered significant hits. Overlaps and unique hits between experimental conditions (parental ± pyruvate, dKO) were assessed using UpSetR v1.4.0. Waterfall plots of beta scores were generated using ggplot2 v3.5.1, with significant hits highlighted.

### Single-Cell RNA Sequencing (scRNA-seq)

Tumors were cut into small fragments and enzymatically digested in DMEM supplemented with 5 mg/mL collagenase type II (ls004176, Worthington) and 50 μg/mL DNase I (D4527-40KU, Sig-ma-Aldrich) for 40 min at 37°C and further processed in gentleMACS C tubes on gentleMACS program (m_impTumor_02). The digested tissue was passed through 70 μm cell strainers to obtain single-cell suspensions, followed by erythrocyte lysis on ice. Cell viability was assessed (>85%). Single-cell suspensions were processed using a droplet-based 31 single-cell RNA-sequencing platform (10x Genomics) Chromium Next GEM Single Cell 3 Gel Bead Kit v3.1, targeting recovery of 6,000 cells per sample. Barcoded cDNA libraries were prepared using single wells of a PCR tube strip, ensuring all cells from each condition were processed with identical reagents in the same master mix and reaction environment. For each experiment, all conditions were handled in parallel using the same thermal cycler. Libraries were sequenced on an Illumina NovaSeq 6000 (See Table S10 for quality metric). Reads were aligned to the mouse (GRCm39) reference genomes using the Cell Ranger pipeline (v8.0.1, 10x Genomics) and downstream analysis was performed with Seurat (Table S10). Gene expression matrices were filtered to retain only genes expressed in at least 50 cells and with a row mean expression ≥ 0.01. Low-quality cells (fewer than 201 detected genes), high-complexity cells (more than 7,500 genes expressed, indicative of potential doublets), and cells with >15% of total unique molecular identifiers (UMIs) mapped to mitochondrial genes were excluded. The remaining gene expression values were normalized and log-transformed using the log1p function in the Seurat package.

The high-throughput sequencing data discussed in this publication have been deposited in NCBI’s Gene Expression Omnibus and will be accessible through GEO Series accession number GSE316786 at www.ncbi.nlm.nih.gov.

## Statistical Analysis

All experiments were repeated at least three times unless stated otherwise. Statistical analysis was performed using GraphPad Prism 10.4.2. Two-tailed unpaired Student’s t-test or ANOVA with post hoc tests were used where appropriate. p < 0.05 was considered statistically signifi-cant.

## Materials Availability

All unique/stable materials generated in this study are available upon request.

## Data and Code Availability

All raw and processed data from metabolomics experiments and CRISPR screen, as well as code used in this study, are available upon request. scRNA-seq data have been deposited in NCBI’s Gene Expression Omnibus and will be accessible through GEO Series accession number GSE316786 at www.ncbi.nlm.nih.gov.

## ACKNOWLEDGEMENTS

We acknowledge the financial support from the Czech Science Foundation (24-10167S, J.R), European Research Council (101042031 – InterMet, K.R.), MSCA (101027977 – MetaCross, K.R.), EMBO (5068, K.R.), MEYS and ERDF/ESF (project CZ.02.01.01/00/22_010/0008608, J.W.S.), MSCA WIDERA (101068637 - EC-InterCom, P.H.), GAUK (1435320, M.M.), MISTI Global Seed Funds (0000000710, K.R. and M.G.V.H.) and by funding from the Czech Academy of Sciences to the Institute of Biotechnology (RVO86652036). M.G.V.H. additionally acknowledges funding from R35CA242379, P30CA014051, the Ludwig Center at MIT, and the MIT Center for Precision Cancer Medicine. S.I.F. is supported by a postdoctoral mandate for fundamental research from the Foundation Against Cancer (Stichting Tegen Kanker, 2023-043). S.-M.F. acknowledges funding from FWO Projects, Fonds Baillet Latour, Francqui Stichting, Foundation ARC, KU Leuven, Stichting tegen Kanker and the Interuniversity BOF (iBOF) programme. J.N. additionally acknowledges funding from the Czech Science Foundation (21-04607X) and Czech Health Research Council (NW25-03-00282).

We used services of the Czech Centre for Phenogenomics at the Institute of Molecular Genetics (IMG) supported by the Czech Academy of Sciences (RVO68378050) and by MEYS CR (LM2023036 Czech Centre for Phenogenomics). Computational resources were provided by the e-INFRA CZ project (ID:90254), supported by the Ministry of Education, Youth and Sports of the Czech Republic. We acknowledge assistance form the Koch Institute/Whitehead Metabolomics Core and the Imaging Methods Core Facility of Faculty of Science, Charles University at BIOCEV.

## AUTHOR CONTRIBUTIONS

Conceptualization: KR, JR; Investigation: M.M., K.D., A.A., P.J., J.W.S., P.H., M.B.M., S.I.F., A.S.-Z., R.St., I.M., Z.C., L.K., J.F.-G., A.B., B.M.; Resources: J.N., S-M.F., M.G.V.H., K.R., J.R.; Funding Ac-quisition: J.N., S.-M.F., M.G.V.H., K.R., J.R.; Writing – Original Draft: M.M., K.R., J.R.; Writing - review & editing: All authors; Supervision: A.B., R.Se., S.-M.F., D.A.T., M.G.V.H., K.R., J.R.

## SUPPLEMENTARY FIGURE LEGENDS

**Figure S1:**
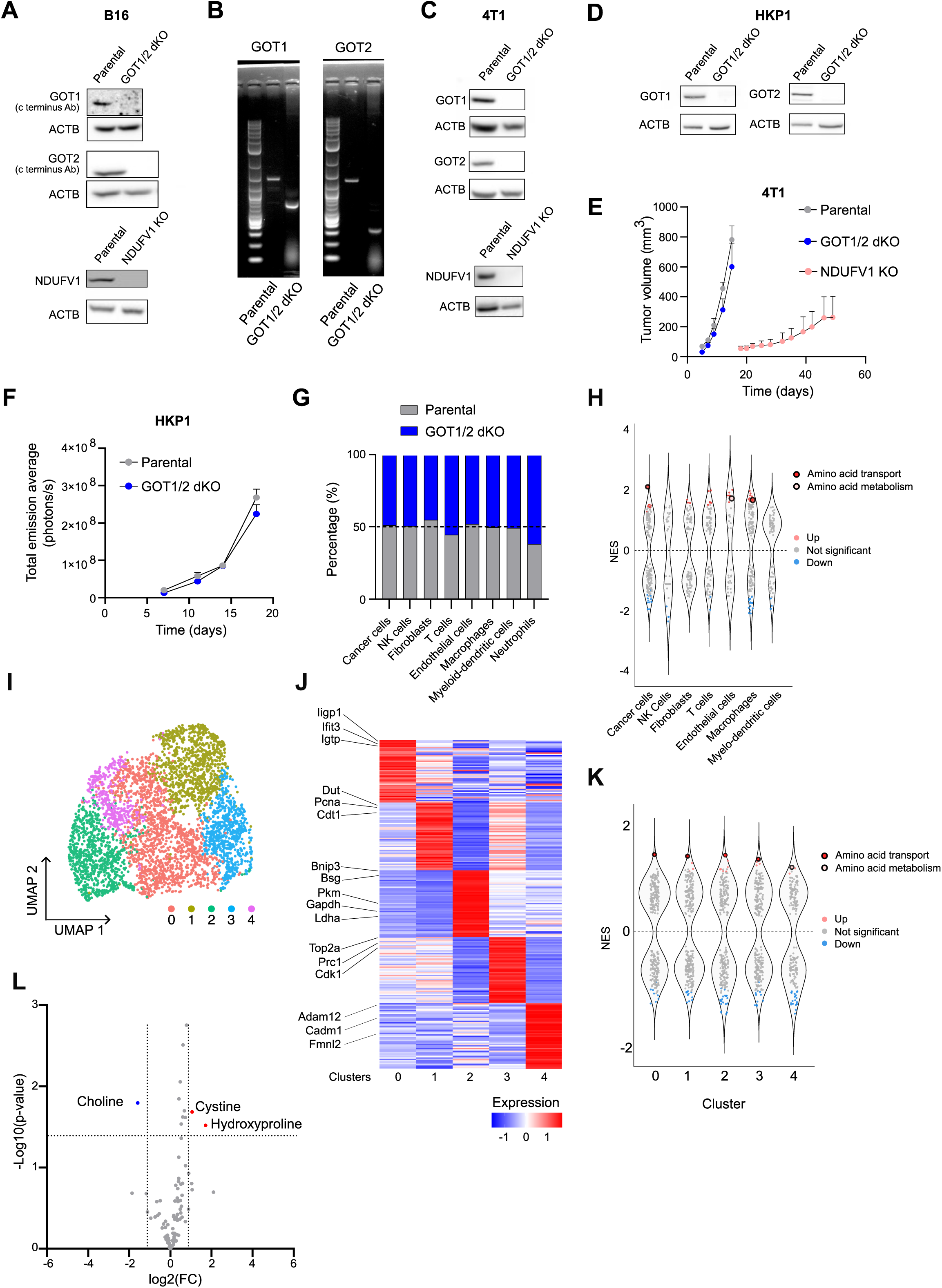
Cellular models and tumor profiling. **A**) Deletion of GOT1 and GOT2 protein in GOT1/2 dKO B16 cells confirmed by C-terminus target-ing antibodies (top) and deletion of NDUFV1 protein in NDUFV1 KO B16 cells, determined by western blot. Representative data from multiple (n≥3) measurements. **B**) Gene editing of GOT1 and GOT2 genes in GOT1/2 dKO B16 cells, determined by DNA gel elec-trophoresis. Representative data from multiple (n≥3) measurements. **C**) Deletion of GOT1, GOT2 (using N-terminal antibodies) and NDUFV1 protein in GOT1/2 dKO and NDUFV1 KO 4T1 cells determined by western blot. Representative data from multiple (n≥3) measurements. **D**) Deletion of GOT1 and GOT2 protein in GOT1/2 dKO HKP1 cells determined by western blot using N-terminal antibodies. Representative data from multiple (n≥3) measurements. **E**) Tumor growth kinetics of orthotopic mouse lung adenocarcinoma tumors generated by tail vein injection of parental and GOT1/2 dKO HKP1 cells (mean ± SEM, n = 5, ∗p < 0.05, two-way ANOVA). **F**) Tumor growth kinetics of mouse mammary carcinoma tumors generated by s.c. injection of parental, GOT1/2 dKO and NDUFV1 KO 4T1 cells (mean ± SEM, n = 5, ∗p < 0.05, two-way ANO-VA). **G**) Relative contribution of parental (grey) and GOT1/2 dKO (blue) tumors to each population determined by scRNAseq. **H**) Differential variation gene set enrichment analysis for metabolic pathways in each cell popu-lation. Each dot represents a metabolic pathway. Amino acid transport and amino acid metabo-lism are highlighted. Note: population of neutrophils showed no significantly changed metabolic pathways. Up, significantly increased in GOT1/2 dKO tumors, down, significantly decreased therein. **I**) UMAP dimensionality reduction plot of single-cell transcriptome of cancer cells from parental and GOT1/2 dKO, color-coded by cluster. **J**) Heatmap of gene expression levels of the top 50 marker genes for clusters of cancer cells from parental and GOT1/2 dKO tumors. Colors represent row-wise scaled gene expression with a mean of 0 and SD of 1 (Z scores) **K**) Differential variation gene set enrichment analysis for metabolic pathways in each cluster of cancer cells. Each dot represents a metabolic pathway. Amino acid transport and amino acid metabolism are highlighted. Up, significantly increased in GOT1/2 dKO tumors; down, signifi-cantly decreased therein. **L**) Quantification of metabolites in plasma of GOT1/2 dKO versus parental B16 tumor bearing mice measured by LC-MS. Blue indicates significantly decreased metabolites and red indicates significantly increased metabolites in plasma of GOT1/2 dKO tumor bearing mice relative to pa-rental B16 tumor bearing mice (n=3, significant hits log2FC > 1, p < 0.05).

**Figure S2:**
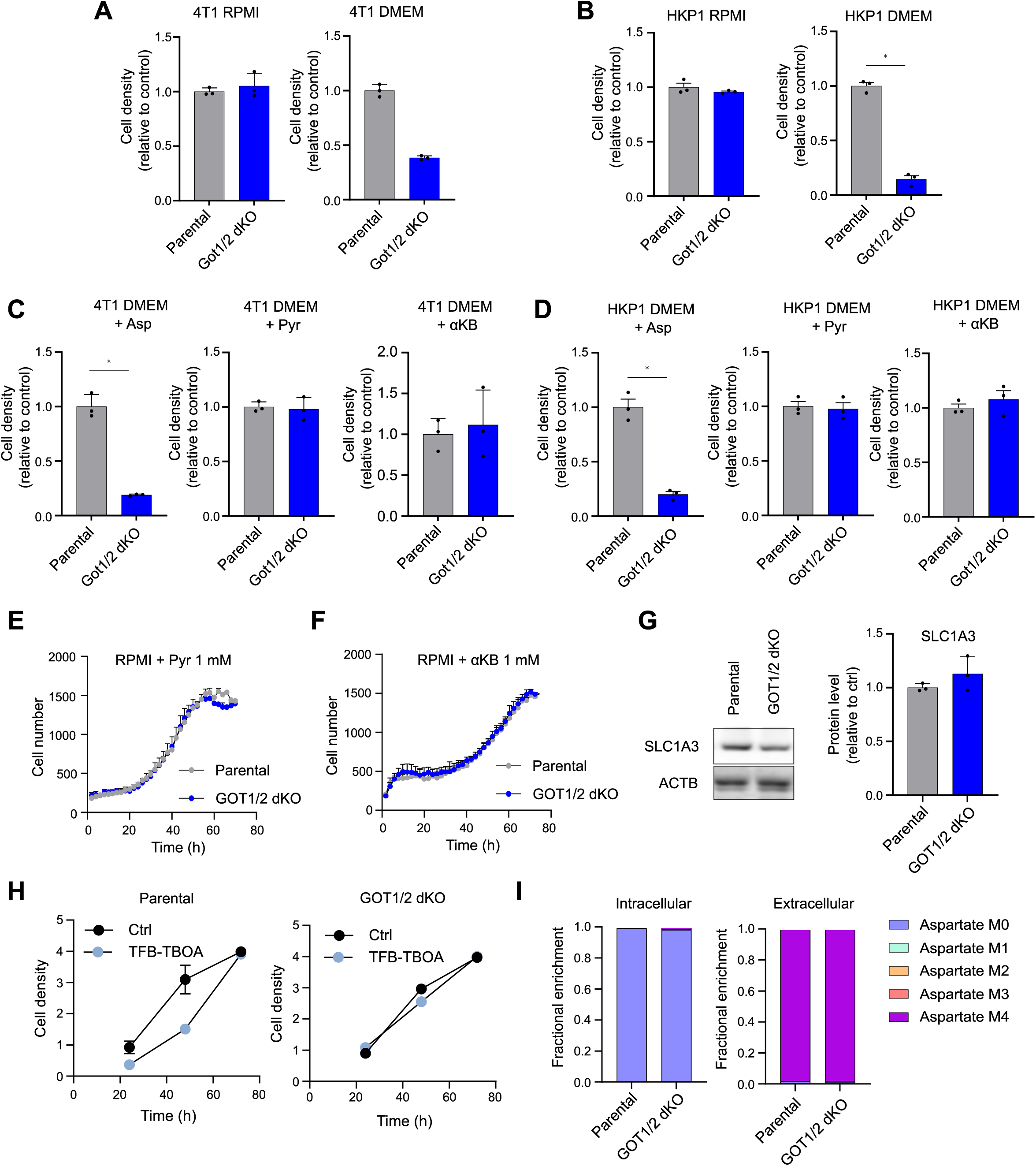
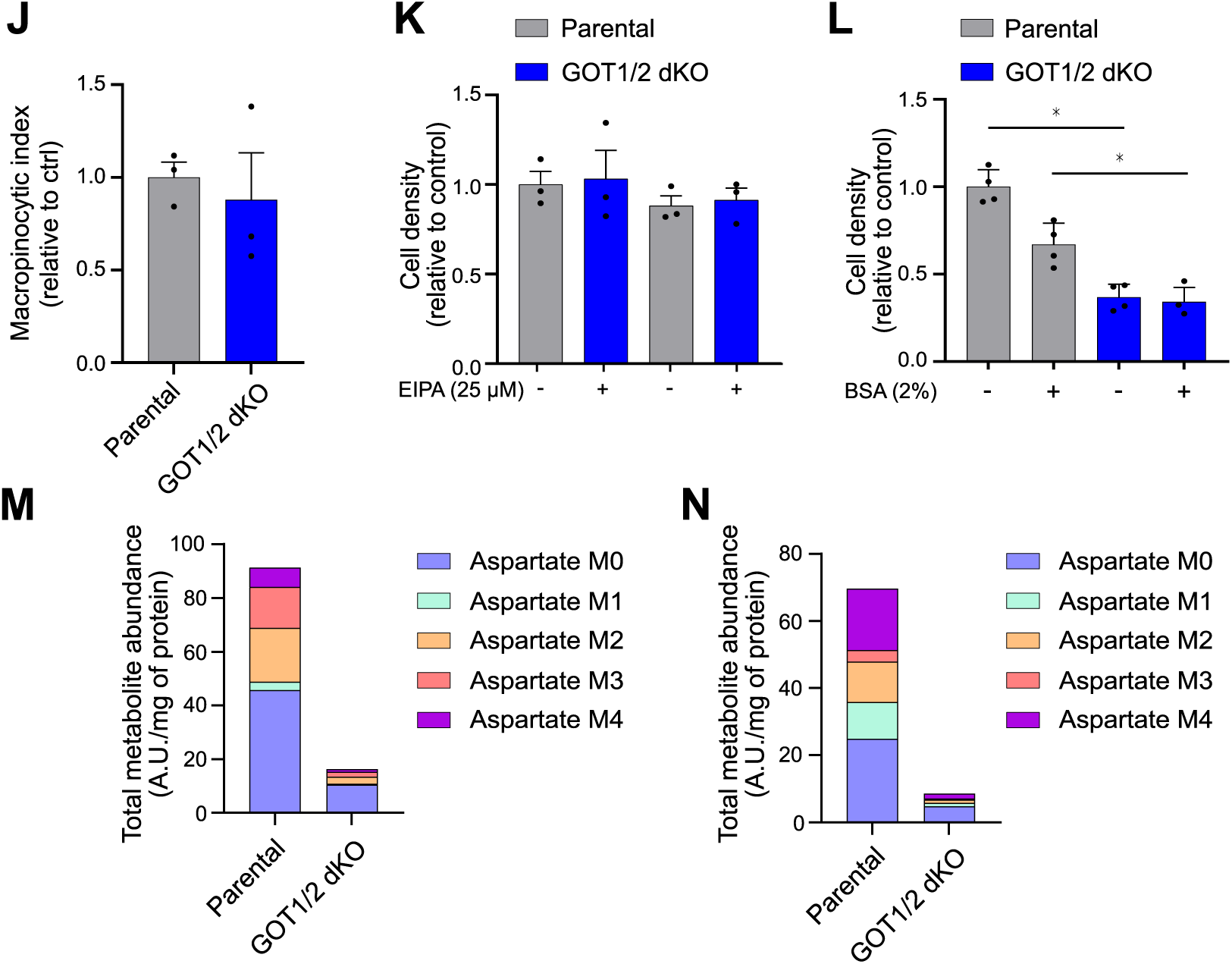
RPMI supplementations and metabolism of aspartate in GOT1/2 dKO. **A**) Proliferation of parental and GOT1/2 dKO 4T1 cells in RPMI or DMEM medium, measured by crystal violet assay after 72 h (mean ± SEM, n = 3, ∗p < 0.05, unpaired two-tailed t test). **B**) Proliferation of parental and GOT1/2 dKO HKP1 cells in RPMI or DMEM medium, measured by crystal violet assay after 72 h (mean ± SEM, n = 3, ∗p < 0.05, unpaired two-tailed t test). **C**) Proliferation of parental and GOT1/2 dKO 4T1 cells in DMEM medium supplemented with 2mM aspartate, 1mM pyruvate or 1mM α-ketobutyrate, measured by crystal violet assay after 72 h (mean ± SEM, n = 3, ∗p < 0.05, unpaired two-tailed t test). **D**) Proliferation of parental and GOT1/2 dKO HKP1 cells in DMEM medium supplemented with 2mM aspartate, 1mM pyruvate or 1mM α-ketobutyrate, measured by crystal violet assay after 72 h (mean ± SEM, n = 3, ∗p < 0.05, unpaired two-tailed t test). **E**) Proliferation of parental and GOT1/2 dKO B16 cells in RPMI medium, supplemented with 1 mM pyruvate, measured by real-time cell proliferation assay (mean ± SEM, n = 3, ∗p < 0.05, un-paired two-tailed t test). **F**) Proliferation of parental and GOT1/2 B16 dKO cells in RPMI medium, supplemented with 1 mM α-ketobutyrate, measured by real-time cell proliferation assay (mean ± SEM, n = 3, ∗p < 0.05, unpaired two-tailed t test). **G**) Representative image (left) and quantification of SLC1A3 protein in parental and GOT1/2 dKO B16 cells determined by western blot. Mean ± SEM, n = 3, ns p > 0.05, unpaired two-tailed t test). **H**) Proliferation of parental (left) and GOT1/2 dKO (right) B16 cells in RPMI medium, in the pres-ence/absence of 1mM TFB-TBOA (SLC1A3 inhibitor), measured by crystal violet assay after 72 h (mean ± SEM, n = 3, ns > 0.05, unpaired two-tailed t test). **I**) Relative C13 exchange in intracellular (left) and extracellular (right) aspartate in parental and GOT1/2 dKO B16 cells cultured in the presence of U-C13 150 µM aspartate in DMEM medium, measured by GC-MS. **J**) Quantification of macropinocytotic index using rhodamine labelled dextran, measured fluo-rescence using microscopy in RPMI medium (mean ± SEM, n = 3, ns > 0.05, unpaired two-tailed t test). **K**) Proliferation of parental and GOT1/2 dKO B16 cells in RPMI medium, in the pres-ence/absence of 25 µM EIPA (micropinocytosis inhibitor), measured by crystal violet assay after 72 h (mean ± SEM, n = 3, ns > 0.05, unpaired two-tailed t test). **L**) Proliferation of parental and GOT1/2 dKO B16 cells in DMEM medium supplemented or not with 2% BSA, measured by crystal violet assay after 72 h (mean ± SEM, n = 3, ns > 0.05, unpaired two-tailed t test). **M**) Relative exchange of U- C into aspartate in parental and GOT1/2 dKO B16 cells, cultured in DMEM medium containing 11 mM U-C13 glucose, measured by GC-MS. **N**) Relative exchange of U- C into aspartate in parental and GOT1/2 dKO B16 cells, cultured in DMEM medium containing 2 mM U-C13 glutamine, measured by GC-MS.

**Figure S3:**
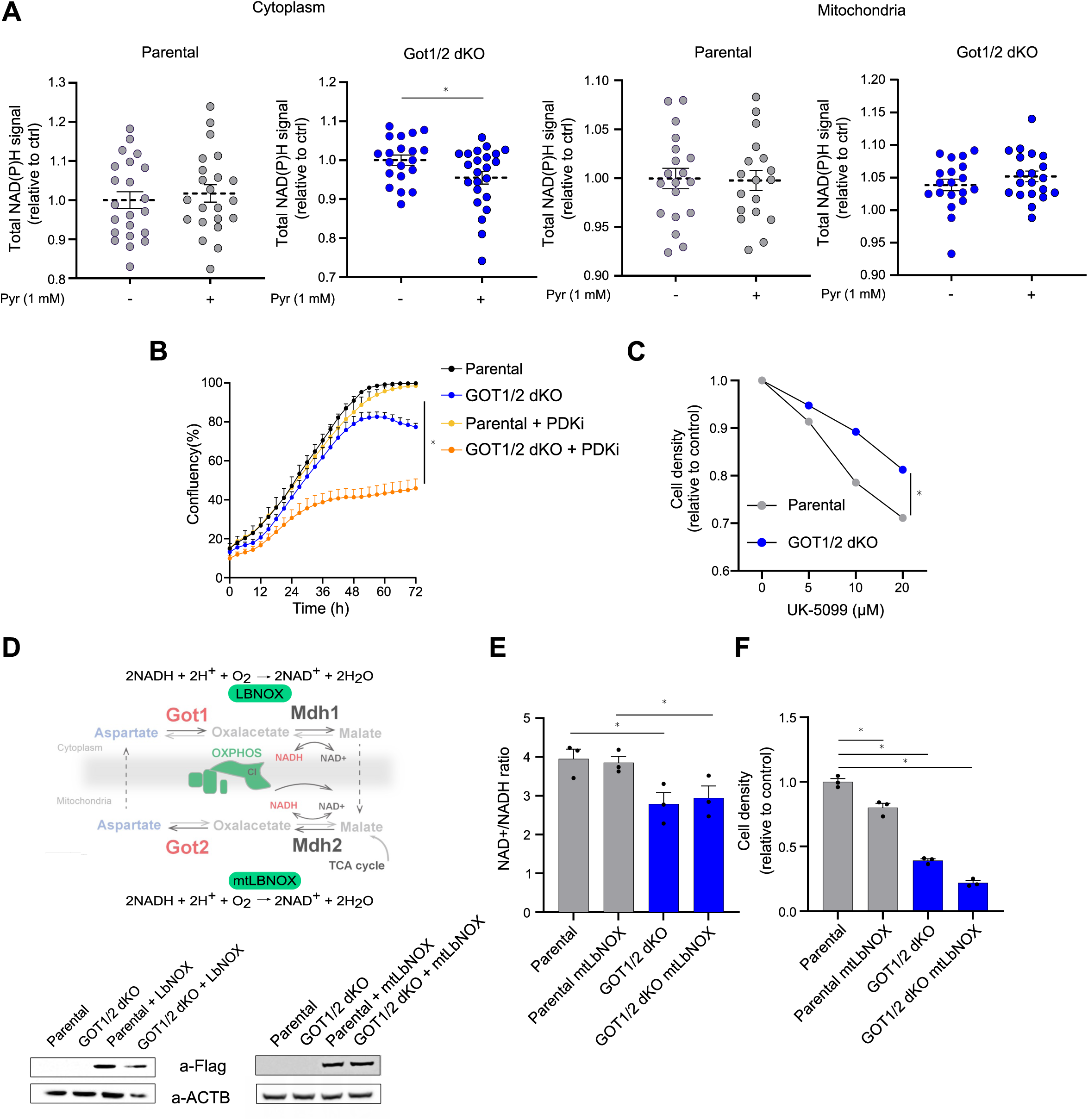
Interventions with cytosolic redox balance. **A**) Total NAD(P)H signal in the cytosol and in mitochondria of parental and GOT1/2 dKO B16 cells cultured in DMEM medium supplemented with 1 mM pyruvate, measured by 2-photon fluorescence-lifetime imaging microscopy (mean ± SEM, n > 19 wells, ∗p < 0.05, unpaired two-tailed t test). Note that the mitochondrial signal comprises also the signal from cytoplasm above and below the mitochondria. **B**) Proliferation of parental and GOT1/2 dKO B16 cells in DMEM medium, supplemented with pyruvate, in the absence/presence of PDK1-4 inhibitor, measured by real-time cell proliferation assay (mean ± SEM, n = 3, ∗p < 0.05, unpaired two-tailed t test). **C**) Proliferation of parental and GOT1/2 dKO B16 cells in the absence or increasing concentra-tion of UK-5099 (inhibitor of mitochondrial pyruvate carrier) grown DMEM medium, supplemented with 1mM pyruvate, measured by Incucyte (mean ± SEM, n = 3, ∗p < 0.05, unpaired two-tailed t test). **D**) A scheme (left) of malate-aspartate shuttle with alternative oxidases (LbNOX) localization and their metabolic reactions. Representative image (right) of flag-tagged LbNOX and mitoLBNOX expressed proteins in parental and GOT1/2 dKO B16 cells determined by western blot. **E**) NAD+/NADH ratio in parental and GOT1/2 dKO B16 cells in the presence/absence of mitoLbNOX expression, grown in DMEM medium, measured by luminescent enzymatic assay (mean ± SEM, n = 3, ∗p < 0.05, unpaired two-tailed t test). **F**) Proliferation of parental and GOT1/2 dKO B16 cells in the presence/absence of mitoLbNOX expression, grown in DMEM medium, measured by crystal violet assay after 72 h (mean ± SEM, n = 3, ns > 0.05, unpaired two-tailed t test).

**Figure S4:**
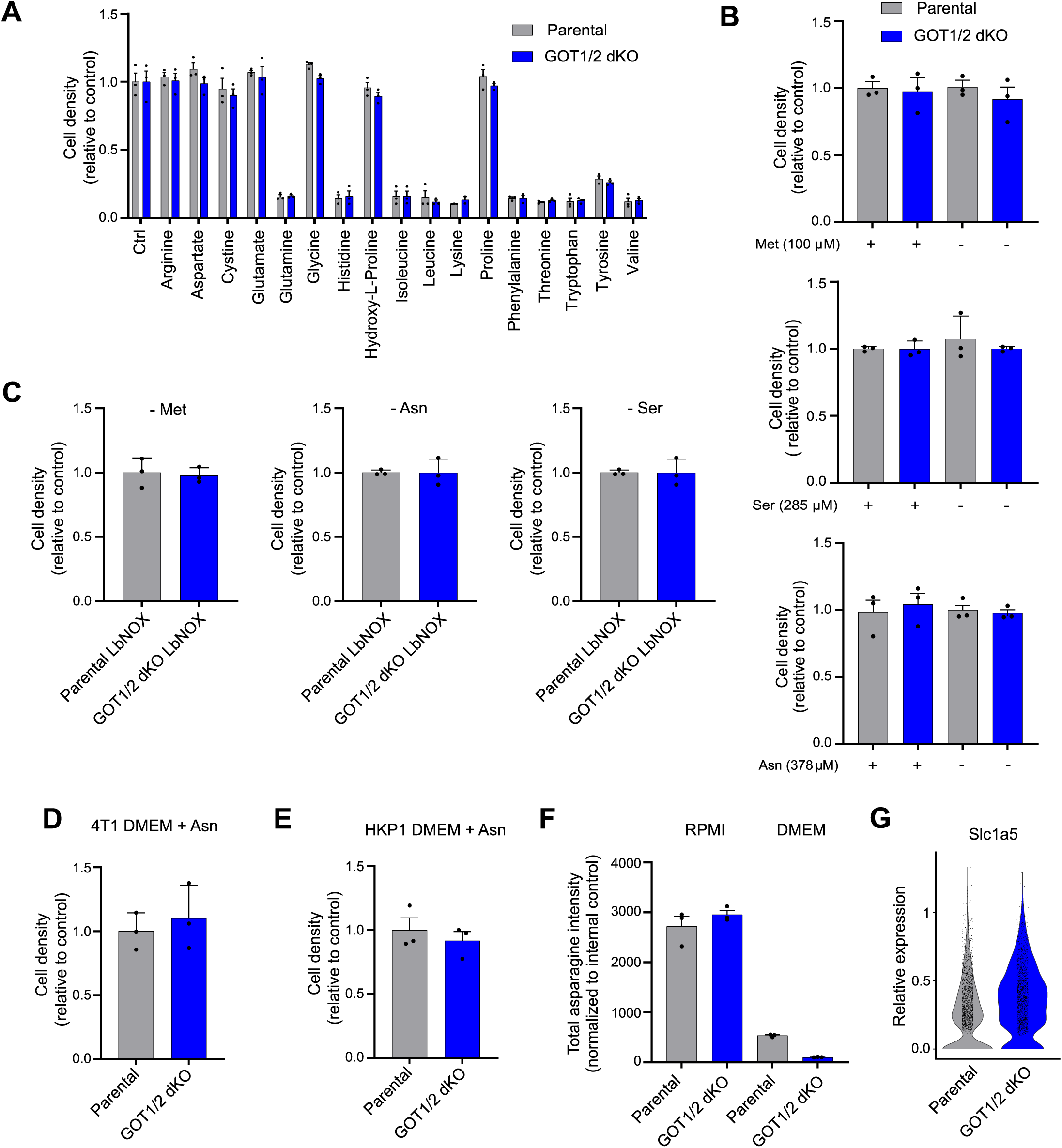
Auxotrophy for amino acids. **A**) Proliferation of parental and GOT1/2 dKO B16 cells grown in reconstituted RPMI medium lacking indicated amino acids, measured by crystal violet assay after 72 h (mean ± SEM, n = 3, ns > 0.05, unpaired two-tailed t test). **B**) Proliferation of parental and GOT1/2 dKO B16 cells in RPMI medium supplemented with 1 mM pyruvate in the presence or absence of methionine, serine or asparagine, measured by crystal violet assay after 72 h (mean ± SEM, n = 3, ns > 0.05, unpaired two-tailed t test). **C**) Proliferation of parental and GOT1/2 dKO B16 cells with or without expression of LbNOX in RPMI medium in the absence of methionine, serine or asparagine, measured by crystal violet assay after 72 h (mean ± SEM, n = 3, ns > 0.05, unpaired two-tailed t test). **D**) Proliferation of parental and GOT1/2 dKO 4T1 cells in DMEM medium supplemented with asparagine (340 µM) measured by crystal violet assay after 72 h (mean ± SEM, n = 3, ∗p < 0.05, unpaired two-tailed t test). **E**) Proliferation of parental and GOT1/2 dKO HKP1 cells in DMEM medium supplemented with asparagine (340 µM) measured by crystal violet assay after 72 h (mean ± SEM, n = 3, ∗p < 0.05, unpaired two-tailed t test). **F**) Intracellular levels of asparagine in parental and GOT1/2 dKO B16 cells grown in RPMI or DMEM medium, measured by GC-MS (mean ± SEM, n = 3, ∗p < 0.05, unpaired two-tailed t test). **G**) Expression of the Slc1a5 transcript in the cancer cell subset of parental and GOT1/2 dKO B16 tumors measured by single-cell RNA sequencing. Violin plots show the distribution of normal-ized transcript expression across individual cells.

**Figure S5:**
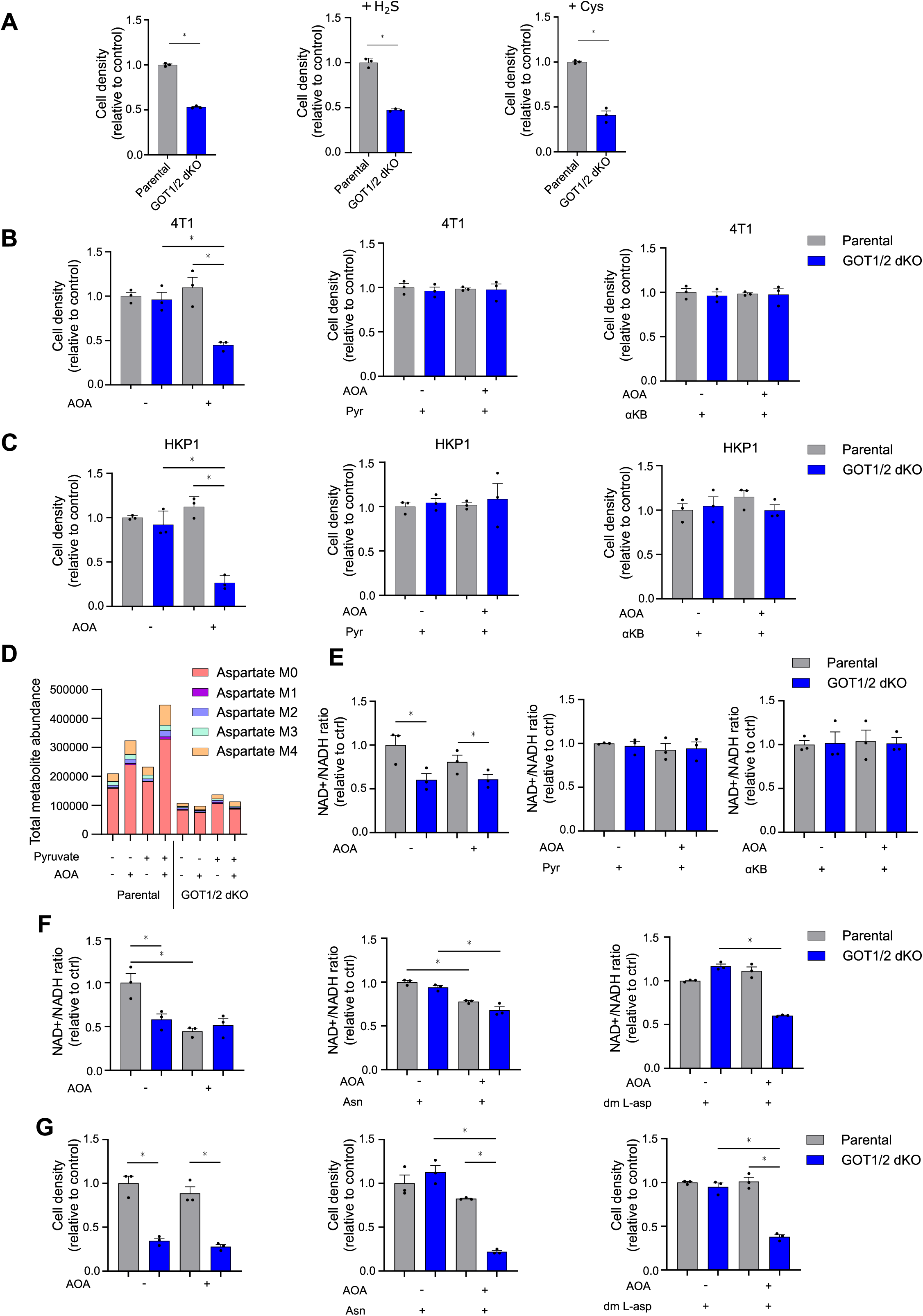
impact of the aminooxyacetic acid treatment. **A**) Proliferation of parental and GOT1/2 dKO B16 cells cultured in DMEM medium supplemented or not with 100 µM H_2_S or with 100 µM cystine, measured by crystal violet assay after 72 h (mean ± SEM, n = 3, ns > 0.05, unpaired two-tailed t test). **B**) Proliferation of parental and GOT1/2 dKO 4T1 cells cultured in RPMI medium in the presence or absence of 300 µM AOA, supplemented with 1mM pyruvate or 1mM α-ketobutyrate, meas-ured by crystal violet assay after 72 h (mean ± SEM, n = 3, ns > 0.05, unpaired two-tailed t test). **C**) Proliferation of parental and GOT1/2 dKO HKP1 cells cultured in RPMI medium in the pres-ence or absence of 300 µM AOA, supplemented with 1mM pyruvate or 1mM α-ketobutyrate, measured by crystal violet assay after 72 h (mean ± SEM, n = 3, ns > 0.05, unpaired two-tailed t test). **D**) Exchange of U-^13^C into aspartate in parental and GOT1/2 dKO B16 cells cultured in DMEM medium supplemented with 2 mM U- C L-glutamine in the presence or absence of 1 mM pyruvate and 300 µM AOA, measured by GC-MS. **E**) NAD^+^/NADH ratio in parental and GOT1/2 dKO B16 cells cultured in DMEM medium supple-mented or not with 1mM pyruvate or 1 mM α-ketobutyrate (α-KB), in the presence or absence of 300 µM AOA, measured by luminescent enzymatic assay (mean ± SEM, n = 3, ∗p < 0.05, unpaired two-tailed t test). **F**) NAD /NADH ratio in parental and GOT1/2 dKO B16 cells cultured in DMEM medium supple-mented or not with 340 µM asparagine or 2 mM dimethyl aspartate (dm-L-asp) in the presence or absence of 300 µM AOA, measured by luminescent enzymatic assay (mean ± SEM, n = 3, ∗p < 0.05, unpaired two-tailed t test). **G**) Proliferation of parental and GOT1/2 dKO B16 cells cultured in DMEM medium supplement-ed or not with 340 µM asparagine or 2 mM dimethyl aspartate (dm-L-asp) in the presence or absence of 300 µM AOA, measured by crystal violet assay after 72 h (mean ± SEM, n = 3, ∗p < 0.05, unpaired two-tailed t test).

**Figure S6:**
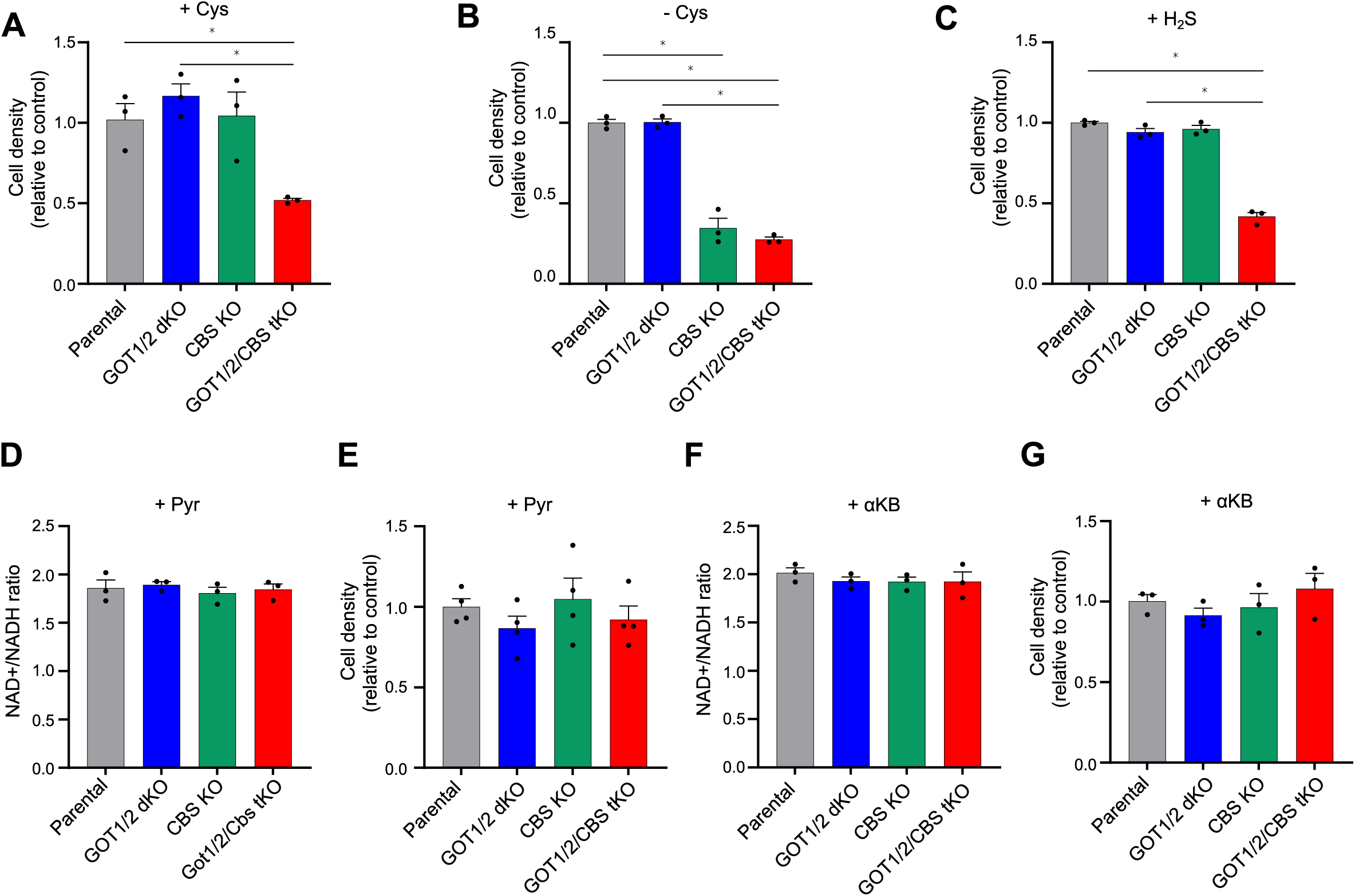
Impact of CBS KO in GOT1/2 dKO cells. **A**) Proliferation of parental GOT1/2 dKO, CBS KO, and GOT1/2/CBS tKO B16 cells, supplemented with 100 µM cystine, grown in RPMI medium, measured by crystal violet assay (mean ± SEM, n = 3, ns > 0.05, unpaired two-tailed t test). **B**) Proliferation of parental GOT1/2 dKO, CBS KO, and GOT1/2/CBS tKO B16 cells in cystine-free RPMI medium measured by crystal violet assay (mean ± SEM, n = 3, ns > 0.05, unpaired two-tailed t test). **C**) Proliferation of parental and GOT1/2 dKO, and GOT1/2/CBS tKO B16 cells, supplemented with 100 µM hydrogen sulfide, grown in DMEM medium, measured by crystal violet assay after 72 h (mean ± SEM, n = 3, ns > 0.05, unpaired two-tailed t test). **D**) NAD+/NADH ratio in parental and GOT1/2 dKO, CBS KO, and GOT1/2/CBS tKO B16 cells grown in RPMI medium supplemented with 1 mM pyruvate, measured by luminescent enzymat-ic assay (mean ± SEM, n > 3, ∗p < 0.05, unpaired two-tailed t test). **E**) Proliferation of parental and GOT1/2 dKO, CBS KO, and GOT1/2/CBS tKO B16 cells in RPMI medium supplemented with 1 mM pyruvate, measured by crystal violet assay after 72 h (mean ± SEM, n = 3, ns > 0.05, unpaired two-tailed t test). **F**) NAD+/NADH ratio in parental and GOT1/2 dKO, CBS KO, and GOT1/2/CBS tKO B16 cells grown in RPMI medium supplemented with 1 mM α-ketobutyrate, measured by luminescent enzymat-ic assay (mean ± SEM, n = 3, ∗p < 0.05, unpaired two-tailed t test). **G**) Proliferation of parental and GOT1/2 dKO, CBS KO, and GOT1/2/CBS tKO B16 cells in RPMI medium supplemented with 1 mM α-ketobutyrate, measured by crystal violet assay after 72 h (mean ± SEM, n = 3, ns > 0.05, unpaired two-tailed t test).

## SUPPLEMENTARY TABLES

Supplementary table 1: Metabolic pathway enrichment score in tumor environment cell types

Supplementary table 2: Metabolic pathways enrichment score in cancer cells

Supplementary table 3: TIF metabolite levels

Supplementary table 4: Plasma metabolite levels

Supplementary table 5: CRISPR screen hits

Supplementary table 6: Amino acid concentrations in custom-made RPMI

Supplementary table 7: Oligonucleotides for generation of GOT1/2 KOs

Supplementary table 8: m/z ratios and retention times for a panel of metabolites meas-ured in TIF and plasma

Supplementary table 9: Primers for CRISPR screen and PCR conditions Supplementary table 10: scRNAseq Quality metrics and parameters of analysis

## REFERENCES

1. Garcia-Bermudez, J. et al. Aspartate is a limiting metabolite for cancer cell proliferation under hypoxia and in tumours. Nat Cell Biol 20, 775–781 (2018).

2. De Falco, P. et al. Hindering NAT8L expression in hepatocellular carcinoma increases cy-tosolic aspartate delivery that fosters pentose phosphate pathway and purine biosynthesis promoting cell proliferation. Redox Biol 59, 102585 (2023).

3. Rabinovich, S. et al. Diversion of aspartate in ASS1-deficient tumours fosters de novo pyrimidine synthesis. Nature 527, 379–383 (2015).

4. Helenius, I.T., Madala, H.R. & Yeh, J.J. An Asp to Strike Out Cancer? Therapeutic Possibili-ties Arising from Aspartate’s Emerging Roles in Cell Proliferation and Survival. Biomolecules 11, 1–11 (2021).

5. Soon, J.W. et al. Aspartate in tumor microenvironment and beyond: Metabolic interac-tions and therapeutic perspectives. Biochim Biophys Acta Mol Basis Dis 1870, 167451 (2024).

6. Birsoy, K. et al. An Essential Role of the Mitochondrial Electron Transport Chain in Cell Proliferation Is to Enable Aspartate Synthesis. Cell 162, 540–551 (2015).

7. Sullivan, L.B. et al. Supporting Aspartate Biosynthesis Is an Essential Function of Respira-tion in Proliferating Cells. Cell 162, 552–563 (2015).

8. Sullivan, L.B. et al. Aspartate is an endogenous metabolic limitation for tumour growth. Nat Cell Biol 20, 782–788 (2018).

9. Bajzikova, M. et al. Reactivation of Dihydroorotate Dehydrogenase-Driven Pyrimidine Biosynthesis Restores Tumor Growth of Respiration-Deficient Cancer Cells. Cell Metab 29, 399–416 e310 (2019).

10. Martinez-Reyes, I. et al. Mitochondrial ubiquinol oxidation is necessary for tumour growth. Nature 585, 288–292 (2020).

11. Kim, W. et al. Polyunsaturated Fatty Acid Desaturation Is a Mechanism for Glycolytic NAD(+) Recycling. Cell Metab 29, 856–870 e857 (2019).

12. Liu, S. et al. Glycerol-3-phosphate biosynthesis regenerates cytosolic NAD(+) to alleviate mitochondrial disease. Cell Metab 33, 1974–1987 e1979 (2021).

13. Tan, A.S. et al. Mitochondrial genome acquisition restores respiratory function and tu-morigenic potential of cancer cells without mitochondrial DNA. Cell Metab 21, 81–94 (2015).

14. Weinberg, F. et al. Mitochondrial metabolism and ROS generation are essential for Kras-mediated tumorigenicity. Proc Natl Acad Sci U S A 107, 8788–8793 (2010).

15. Balsa, E. et al. Defective NADPH production in mitochondrial disease complex I causes inflammation and cell death. Nat Commun 11, 2714 (2020).

16. Krall, A.S. et al. Asparagine couples mitochondrial respiration to ATF4 activity and tumor growth. Cell Metab 33, 1013–1026 e1016 (2021).

17. Garcia-Bermudez, J. et al. Adaptive stimulation of macropinocytosis overcomes aspar-tate limitation in cancer cells under hypoxia. Nat Metab 4, 724–738 (2022).

18. Agalioti, T. et al. Mutant KRAS promotes malignant pleural effusion formation. Nat Commun 8, 15205 (2017).

19. Hart, M.L. et al. Succinate dehydrogenase loss suppresses pyrimidine biosynthesis via succinate-mediated inhibition of aspartate transcarbamylase. Nat Metab 8, 1390–1409 (2026).

20. Kerk, S.A. et al. Metabolic requirement for GOT2 in pancreatic cancer depends on envi-ronmental context. Elife 11, e73245–e73245 (2022).

21. Titov, D.V. et al. Complementation of mitochondrial electron transport chain by manipu-lation of the NAD+/NADH ratio. Science 352, 231–235 (2016).

22. Brunner, J.S. et al. Aspartate availability drives differential engagement of the malate-aspartate shuttle. Mol Cell 86, 954–967 e957 (2026).

23. Zhu, X.G. et al. Functional Genomics In Vivo Reveal Metabolic Dependencies of Pancreat-ic Cancer Cells. Cell Metab 33, 211–221 e216 (2021).

24. Werge, M.P. et al. The Role of the Transsulfuration Pathway in Non-Alcoholic Fatty Liver Disease. J Clin Med 10, 1081 (2021).

25. Krall, A.S., Xu, S., Graeber, T.G., Braas, D. & Christofk, H.R. Asparagine promotes cancer cell proliferation through use as an amino acid exchange factor. Nat Commun 7, 11457 (2016).

26. Carroll, W.R., Stacy, G.W. & Du Vigneaud, V. alpha-Ketobutyric acid as a product in the enzymatic cleavage of cystathionine. J Biol Chem 180, 375–382 (1949).

27. Lesner, N.P., Gokhale, A.S., Kota, K., DeBerardinis, R.J. & Mishra, P. alpha-ketobutyrate links alterations in cystine metabolism to glucose oxidation in mtDNA mutant cells. Metab Eng 60, 157–167 (2020).

28. Rosalki, S.B. & Wilkinson, J.H. Reduction of alpha-ketobutyrate by human serum. Nature 188, 1110–1111 (1960).

29. Liang, J., Han, Q., Tan, Y., Ding, H. & Li, J. Current Advances on Structure-Function Rela-tionships of Pyridoxal 5’-Phosphate-Dependent Enzymes. Front Mol Biosci 6, 4 (2019).

30. Ascencao, K. & Szabo, C. Emerging roles of cystathionine beta-synthase in various forms of cancer. Redox Biol 53, 102331 (2022).

31. Pavlova, N.N. et al. As Extracellular Glutamine Levels Decline, Asparagine Becomes an Essential Amino Acid. Cell Metab 27, 428–438 e425 (2018).

32. Zhang, J. et al. Asparagine plays a critical role in regulating cellular adaptation to gluta-mine depletion. Mol Cell 56, 205–218 (2014).

33. Halbrook, C.J. et al. Differential integrated stress response and asparagine production drive symbiosis and therapy resistance of pancreatic adenocarcinoma cells. Nat Cancer 3, 1386–1403 (2022).

34. Diehl, F.F., Lewis, C.A., Fiske, B.P. & Vander Heiden, M.G. Cellular redox state constrains serine synthesis and nucleotide production to impact cell proliferation. Nat Metab 1, 861–867 (2019).

35. Broeks, M.H., et al. The malate-aspartate shuttle is important for de novo serine biosyn-thesis. Cell Rep 42, 113043 (2023).

36. Zhu, J. et al. Transsulfuration Activity Can Support Cell Growth upon Extracellular Cyste-ine Limitation. Cell Metab 30, 865–876 e865 (2019).

37. Khan, N.A. et al. mTORC1 Regulates Mitochondrial Integrated Stress Response and Mito-chondrial Myopathy Progression. Cell Metab 26, 419–428 e415 (2017).

38. Hart, M.L. et al. Mitochondrial redox adaptations enable alternative aspartate synthesis in SDH-deficient cells. Elife 12, e78654 (2023).

39. Yap, T.A. et al. Complex I inhibitor of oxidative phosphorylation in advanced solid tumors and acute myeloid leukemia: phase I trials. Nat Med 29, 115–126 (2023).

40. Magalhaes-Novais, S. et al. Mitochondrial respiration supports autophagy to provide stress resistance during quiescence. Autophagy 18, 2409–2426 (2022).

41. Sullivan, M.R., Lewis, C.A. & Muir, A. Isolation and Quantification of Metabolite Levels in Murine Tumor Interstitial Fluid by LC/MS. Bio Protoc 9, e3427 (2019).

42. Abbott, K.L. et al. Metabolite profiling of human renal cell carcinoma reveals tissue-origin dominance in nutrient availability. Elife 13, RP95652 (2024).

43. Kolarova, L. et al. Clusters of Monoisotopic Elements for Calibration in (TOF) Mass Spec-trometry. J Am Soc Mass Spectrom 28, 419–427 (2017).

44. Strohalm, M., Kavan, D., Novak, P., Volny, M. & Havlicek, V. mMass 3: a cross-platform software environment for precise analysis of mass spectrometric data. Anal Chem 82, 4648–4651 (2010).

45. Chong, J., Wishart, D.S. & Xia, J. Using MetaboAnalyst 4.0 for Comprehensive and Inte-grative Metabolomics Data Analysis. Curr Protoc Bioinformatics 68, e86 (2019).

46. Ewels, P., Magnusson, M., Lundin, S. & Kaller, M. MultiQC: summarize analysis results for multiple tools and samples in a single report. Bioinformatics 32, 3047–3048 (2016).

47. Li, W. et al. MAGeCK enables robust identification of essential genes from genome-scale CRISPR/Cas9 knockout screens. Genome Biol 15, 554 (2014).

48. Li, W. et al. Quality control, modeling, and visualization of CRISPR screens with MAGeCK-VISPR. Genome Biol 16, 281 (2015).

49. Satija, R., Farrell, J.A., Gennert, D., Schier, A.F. & Regev, A. Spatial reconstruction of sin-gle-cell gene expression data. Nat Biotechnol 33, 495–502 (2015).

